# Stabilized COre gene and Pathway Election uncovers pan-cancer shared pathways and a cancer specific driver

**DOI:** 10.1101/2021.12.21.473727

**Authors:** Pathum Kossinna, Weijia Cai, Xuewen Lu, Carrie S Shemanko, Qingrun Zhang

## Abstract

Approaches systematically characterizing interactions via transcriptomic data usually follow two systems: (1) co-expression network analyses focusing on correlations between genes; (2) linear regressions (usually regularized) to select multiple genes jointly. Both suffer from the problem of stability: a slight change of parameterization or dataset could lead to dramatic alternations of outcomes. Here, we propose **S**tabilized **Co**re gene and **P**athway **E**lection, or *SCOPE*, a tool integrating bootstrapped LASSO and co-expression analysis, leading to robust outcomes insensitive to variations in data. By applying SCOPE to six cancer expression datasets (BRCA, COAD, KIRC, LUAD, PRAD and THCA) in The Cancer Genome Atlas, we identified core genes capturing interaction effects in crucial pan-cancer pathways related to genome instability and DNA damage response. Moreover, we highlighted the pivotal role of CD63 as an oncogenic driver and a potential therapeutic target in kidney cancer. SCOPE enables stabilized investigations towards complex interactions using transcriptome data.

**Availability:** https://github.com/QingrunZhangLab/SCOPE

## Introduction

Understanding the process of pathogenesis and discovering new therapeutic targets requires discovery of the underlying driver genes in relevant pathways^1–3^. However, determination of the “driver” role of a gene through experimental investigation has only been possible for a handful of genes due to the time consuming and expensive nature of such experiments. Thus, *in-silico* analysis to narrow down candidates of potential genes is vital. Current methods of identifying driver genes involve multi-omics data^4^, and often utilise known biological pathways^5^. Among multi-omics data, transcriptome, i.e., gene expression data, play a pivotal role in biological processes; and are the most available form of -omics data for many diseases including cancers. As such, analyzing transcriptomic data is usually the first step in -omics-directed characterization of diseases.

In practice, selecting differentially expressed (DE) genes by contrasting expression levels in disease and control tissues has been broadly used for the exploration of biological mechanisms of various diseases. Largely due to its simplicity, single-gene-based DE analysis is the most popular method adapted by many researchers^6^. From the perspective of systems biology, it is natural to expect that advanced models analyzing multiple genes jointly should lead to additional in-depth understanding of disease pathology.

Unfortunately, instability of such complex models involving multiple genes appears to be a serious problem; in many situations current methods do not generate consistent results. For instance, in a typical co-expression network-based analysis, a gene-network is built with its nodes representing genes and edges based on their co-expression. The genes that are highly connected with other genes in the network, called “hub” genes, are expected to be important in pathology^7^. As such, many pipelines discovering driver genes incorporate information from co-expression networks and these hub genes into the next phase of multiomics approaches^8–10^. It has, however, been noted that hub genes are not stable, and they are not guaranteed to be driver genes^11^.

Regularized multiple regression methods, which optimize an objective function by adding a regularization term to a likelihood, are widely used in many domains^12–14^ including biomarker selection using transcriptomic data. LASSO (Least Absolute Shrinkage and Selection Operator) and ridge regression are two representative methods of this nature^15,16^. The choice of regularization plays a significant role in the information supplied by the final model: ridge regression will lead to a model containing a large number of genes^15^, which may confer a high predictive power at the cost of little meaningful information for functional characterization; LASSO, in contrast, retains fewer genes^16^, however is inherently unstable in the presence of highly correlated variables^17,18^, which is unfortunately the case of transcriptome data. Indeed, while a logistic LASSO regression can usually identify significant variables in determining case and control, it also tends to provide completely different outcomes with different parameterizations, or even by running a similarly parameterized model multiple times over slightly different data^17,19^. That is why historically, as far as we understand, there have been few efforts using such feature selection methods in identifying underlying genes from transcriptomic data.

Co-expression network analyses and regularized multiple regressions form disconnected fields, which are by themselves unable to produce stable results offering insights into disease pathology. We propose the **S**tabilised **CO**re gene and **P**athway **E**lection, or **SCOPE**, a new tool to reliably discover candidate genes and pathways using transcriptomic data. The new framework represents a synergy between co-expression network analysis and regularized multiple regressions, with two layers of stabilization integrated.

As a proof-of-concept, we applied SCOPE to six cancer datasets from The Cancer Genome Atlas (TCGA)^20,21^ (BRCA, COAD, KIRC, LUAD, PRAD and THCA with 1,041-111, 480-70, 483-54, 387-37, 458-50, 444-53 “Primary Tumor”-“Normal Tissue” samples respectively) to identify novel core genes and their related pathways. Thorough comparisons were carried out against standard methods including LASSO (for the selection step only) and differential expression (DE) analysis as well as differential co-expression (DiffCoEx) analysis. As expected, the core genes selected by SCOPE-Stabilized LASSO indeed are stable with respect to small changes of the input datasets. Despite being significantly fewer than the typical set of genes identified by a standard LASSO, the SCOPE-identified core genes remain highly predictive. Notably, as another line of evidence at the pathway level, pathways identified by SCOPE show significant within- and pan-cancer overlap. In contrast, standard differential co-expression analysis led to significantly lower overlaps across and within cancers. Moreover, as a confirmation by employing connectivity analysis, we found that the core genes indeed play central roles in the shared pathways.

To discover novel insights into cancer pathology based on the stably identified core genes and pathways, we have carefully annotated the data based on our insights into cancers and from the literature. Notably, we shed light on the critical role of *CD63* in the Von Hippel-Lindau tumor suppressor (*VHL*) - hypoxia-inducible factor 1 subunit α (*HIF1A*)/hypoxia-inducible factor 2α (*HIF2A*) - vascular endothelial growth factor A (*VEGFA*) protein (*VHL*-*HIF*-*VEGF*/*VEGFR*) axis^22^, a putative key driver that governs tumorigenesis in kidney cancer. These discoveries may provide interesting insights into the mechanism of cancer, identifying concrete targets for further experimental follow-ups.

## Results

### The SCOPE Framework

SCOPE begins by conducting multiple regression by utilizing a stabilized extension of LASSO, here termed SCOPE-Stabilized LASSO which employs a bootstrap of multiple LASSO models (**Figure 1A** and **B**), leading to a handful of core genes robust to statistical instability (**Figure 1C**). These core genes differ from hub genes in that they are not identified due to their interconnectedness – but due to their power in prediction, whilst still being stable across random samples. These core genes are then used as seed genes for further co-expression analysis (**Figure 1D**). The outcomes of the co-expression analysis of each core gene are then piped into pathway enrichment analysis (**Figure 1E**). The pathways learned from each core gene are finally intersected to provide another level of stabilization (**Figure 1E**). A high-level pseudo-code is included in **Figure 1F**, and the detailed algorithms and design considerations are provided in **Methods** and **Supplementary Materials**. This framework incorporates both optimizations brought by multiple regression and gene-gene interactions identified by co-expression analysis, whilst retaining stability in large part due to two levels of stabilization.

**Figure 1.**
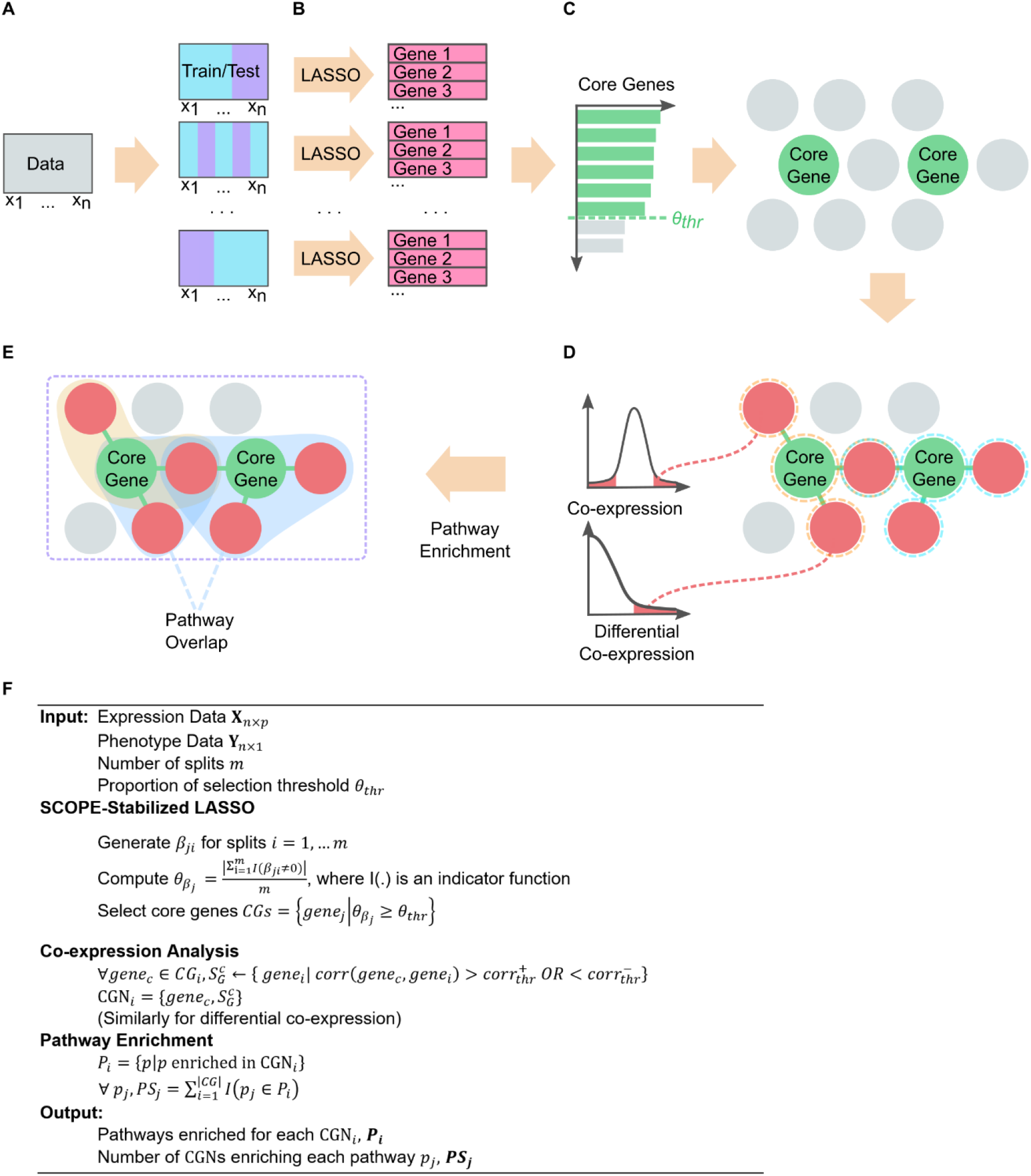
Overview of SCOPE Framework: **A)** Expression data are split multiple times randomly into typical training/test splits with a consistent phenotype ratio as in the original data. **B)** LASSO models are trained on each of the splits and the genes selected in each split are recorded. **C)** Selected genes are ordered by the frequency of occurrence in the splits. Based on a cut-off *θ*_*thr*_ (dotted green line), core genes are identified and are used to identify the Core Gene Networks (CGNs) in (d). **D)** CGNs are identified, indicated by genes circled in orange and blue dashed lines. Null distributions of both differential co-expression and co-expression are used to identify genes significantly interacting with the identified core genes. **E)** Pathway enrichment analysis is conducted for each CGN, and the overlap between CGN-directed pathways will be identified. Finally, substantially overlapped pathways and core genes will be the output. **F)** The algorithm in a simplified high-level pseudo-code. The full version of the algorithm is presented in **Supplementary Materials**.

### SCOPE stably selected considerably fewer core genes while retaining predictive power

SCOPE-Stabilized LASSO identified significantly fewer genes among different splits of the data (**Figure 2A** vertical red dotted lines). The consistently selected genes are named in **Figure 2A** with details in **Supplementary Table S1**. In contrast, on the same data, genes selected by a standard LASSO are much more numerous and vary widely from around 10 to 45 genes (**Figure 2A** colored distributions), documenting the instability of gene selection by standard LASSO.

**Figure 2.**
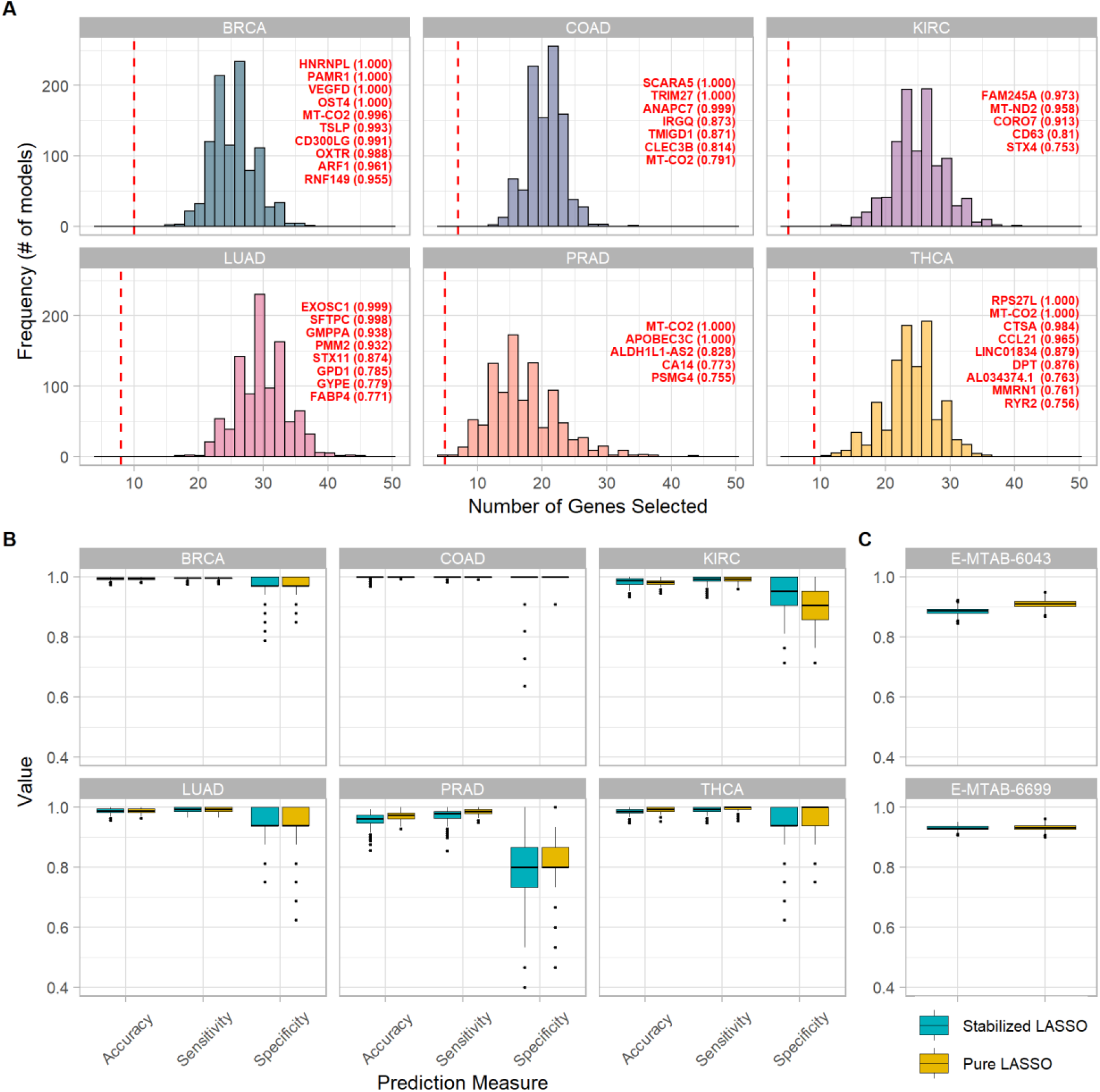
Comparison of SCOPE to Standard LASSO in stability and predictive accuracy. **A)** Histograms of the number of genes selected by standard LASSO (colored distributions) in comparison to SCOPE (vertical red dotted lines) for each cancer. The thresholds chosen for SCOPE-selected core genes were varied: *θthr* = 0.90 for BRCA and *θthr* = 0.75 for KIRC, LUAD, COAD, PRAD and THCA. These thresholds resulted in 5-10 core genes being identified per cancer, identified in the 6 panels for each cancer. **B)** Prediction metrics for SCOPE (core genes) in comparison to standard LASSO in terms of accuracy = (TP+TN)/(TP+TN+FP+FN), sensitivity and specificity. **C)** Prediction accuracy = (TP+TN)/(TP+TN+FP+FN) for two independent micro-array datasets for lung cancer were obtained. In the case of SCOPE, the same core genes identified and indicated in **Supplementary Table S1** were used. For standard LASSO, multiple sets of genes selected by independent LASSO runs in the TCGA LUAD dataset were used to assess the varied distribution (due to the instability).

Despite being much smaller in number, the predictive power of SCOPE selected core genes is close to that obtained by standard LASSO. In-sample validation shows that the few genes identified by SCOPE confer almost the same, and sometimes even higher, predictive power compared with the many genes selected by standard LASSO (**Figure 2B**). We also resorted to external data validation using two micro-array datasets^23,24^ based on the set of core genes identified by SCOPE-Stabilized LASSO (listed in **Supplementary Table S1**) and the multiple runs of standard LASSO (mirroring in practice the range different people might achieve based on the genes they ended up identifying). Evidently, the predictive accuracy remains close to one using standard LASSO (**Figure 2C**). Internal and external validation highlighted the ability of SCOPE-Stabilized LASSO to identify a highly predictive handful of genes that are comparable in prediction accuracy to the many-folds larger number of genes selected by a standard LASSO model. The small margin also indicates that, while models including more genes may be slightly more predictive, they may not all be vital to tumorigenesis. Furthermore, such a large number of genes could be extremely costly to experimentally validate and thus are not an ideal outcome of *in-silico* methods, a problem relieved by SCOPE-Stabilized selection.

The consistency and stability of SCOPE-Stabilized LASSO over standard LASSO was demonstrated by looking at the replicability over multiple runs on the same data (with different splits of training/testing samples). The comparison was conducted by randomly splitting the data into training and testing samples 100 times, using a different random seed for each split. This reflects the effect of choosing a different training sample in a typical usage scenario. Proportions of runs in which genes were selected are shown in **Supplementary Table S2** illustrating the high level of stability obtained by SCOPE-Stabilized LASSO over standard LASSO.

### SCOPE identified pan-cancer pathways, focusing on DNA replication and repair

Via standard co-expression analysis of gene networks, the core genes selected by the SCOPE-Stabilized LASSO were used to form their corresponding Core Gene Networks (CGNs), which in turn were used to identify pathways based on pathway enrichment analysis (**Methods**). The pathways that are identified by multiple CGNs are the output of SCOPE (**Supplementary Table S3**). Many of these pathways fall into the categories of “Cell Growth and Death”, “Replication and Repair” and “Folding, sorting and degradation”. These pathways are highly related to cancer cell immortality and cancer genome damage response. A similar protocol was also conducted using Differential Co-Expression (DiffCoEx) analysis by looking at pathways identified by multiple modules (**Supplementary Table S4**).

To further assess the stability of SCOPE, we analyzed the sharing of core genes between cancers. At the gene level, *MT-CO2* was identified as a core gene by SCOPE in four out of the six cancers. This gene produces the *Cytochrome c oxidase subunit 2* protein which is essential in a mitochondrial process associated with oxidative phosphorylation. Besides *MT-CO2*, no other core gene is shared among cancers, indicating that different cancers may have different core genes if one does not look at higher levels such as pathways.

We then characterized the sharing of pathways across cancers. To quantify the extent of sharing, we first formed a within-cancer statistic *π*_*cancer*_, which denotes the proportion of genes contained in CGNs out of the total number of genes in each pathway. Then, the Pathway Overlap Score, or POS, was calculated as the summation of the *π*_*cancer*_ values over all six cancers. Higher values intuitively indicate higher overlap of the pathway across cancers. By contrasting the results of the three tools, namely SCOPE, DE and DiffCoEx, despite their substantial sharing in terms of pathways identified (**Figure 3A**), showed quite different landscapes in terms of sharing between cancers. Evidently, SCOPE identified both cancer-specific and pan-cancer pathways characterized by its two-spike distribution of POS: the cluster at the low-POS end stands for cancer-specific, while the cluster at the high-POS end indicates pan-caner pathways (**Figure 3B**). In contrast, DE and DiffCoEx distributions both have only one spike at the low-POS spectrum (**Figure 3C and D**). These POS distributions are further detailed in **Supplementary Table S5** for SCOPE, DiffCoEx, and DE, evidencing the potential drawback of DiffCoEx and DE in their inability to repetitively identify key pathways that could be universally vital in cancers. This distinction between SCOPE and standard methods suggests that SCOPE’s stability in discovering pathways can reveal key pathways even in different cancers.

**Figure 3.**
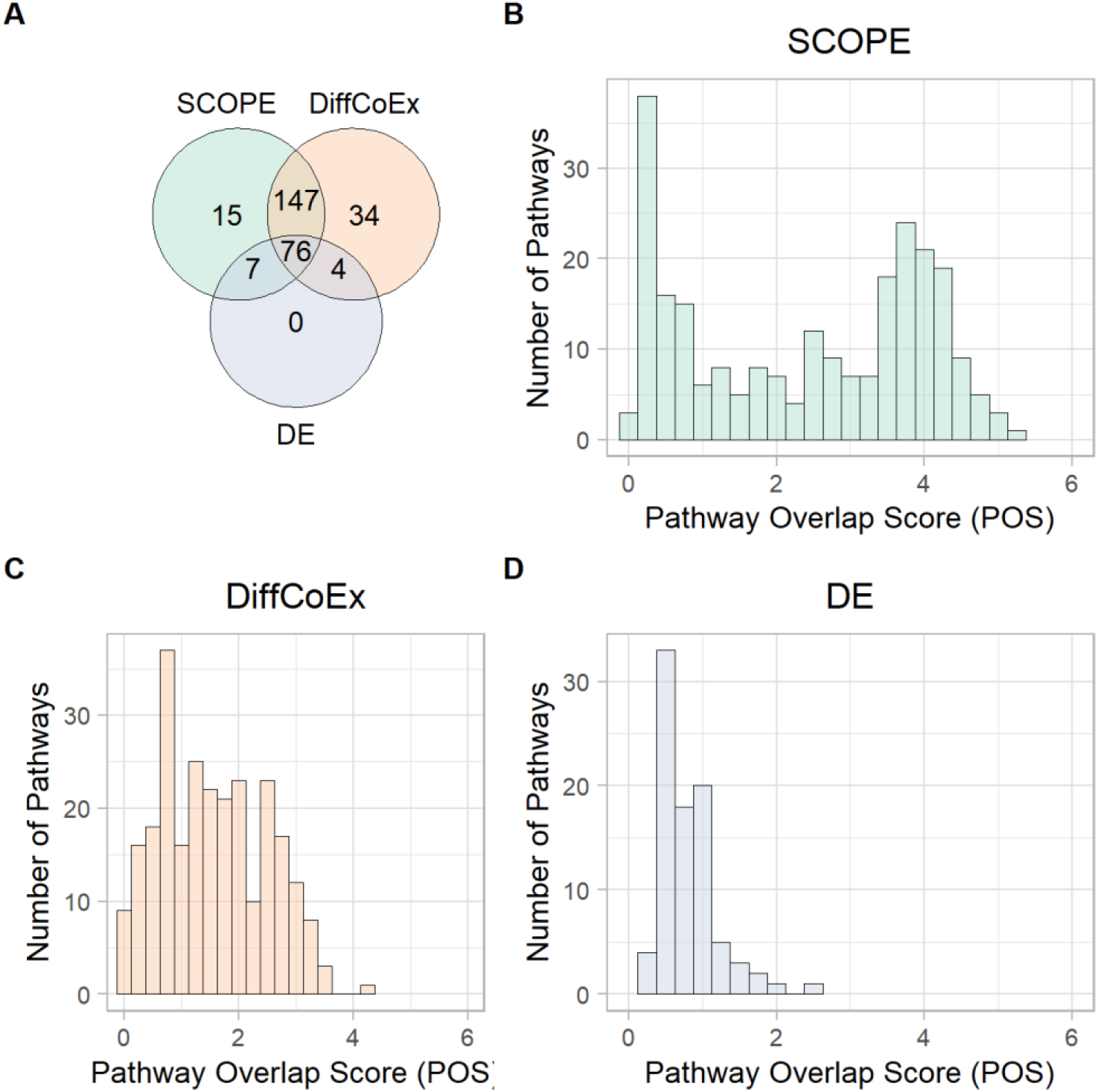
Comparison of SCOPE to alternative methods on pathway identifications: **A)** Pathways identified by DiffCoEx, DE, and the SCOPE are compared for uniqueness and sharing. **B), C) and D)** The Pathway Overlap Score (POS) which indicates the level of enrichment of a pathway across multiple cancers is contrasted amongst the three methods. **B)** SCOPE uncovers both cancer specific (notable by the spike in lower POS), as well as pan-cancer shared pathways (notable by the spike in higher POS), while DE **(C)** and DiffCoEx **(D)** both appear to be more cancer specific than SCOPE as evidenced by the lower distribution of POS.

We then annotated pathways that were identified by SCOPE to check if they were relevant. Investigating pathways related to the hallmarks of cancer^25^ and the proportion of genes in each of these pathways by each of the three methods (**Figure 4A – F**) reveals that SCOPE identifies these hallmarks across cancers quite significantly. **Figure 4G** looks at the POS for these hallmark pathways across the different cancers. SCOPE stands out by identifying the highest proportion of genes involved in these pathways.

**Figure 4.**
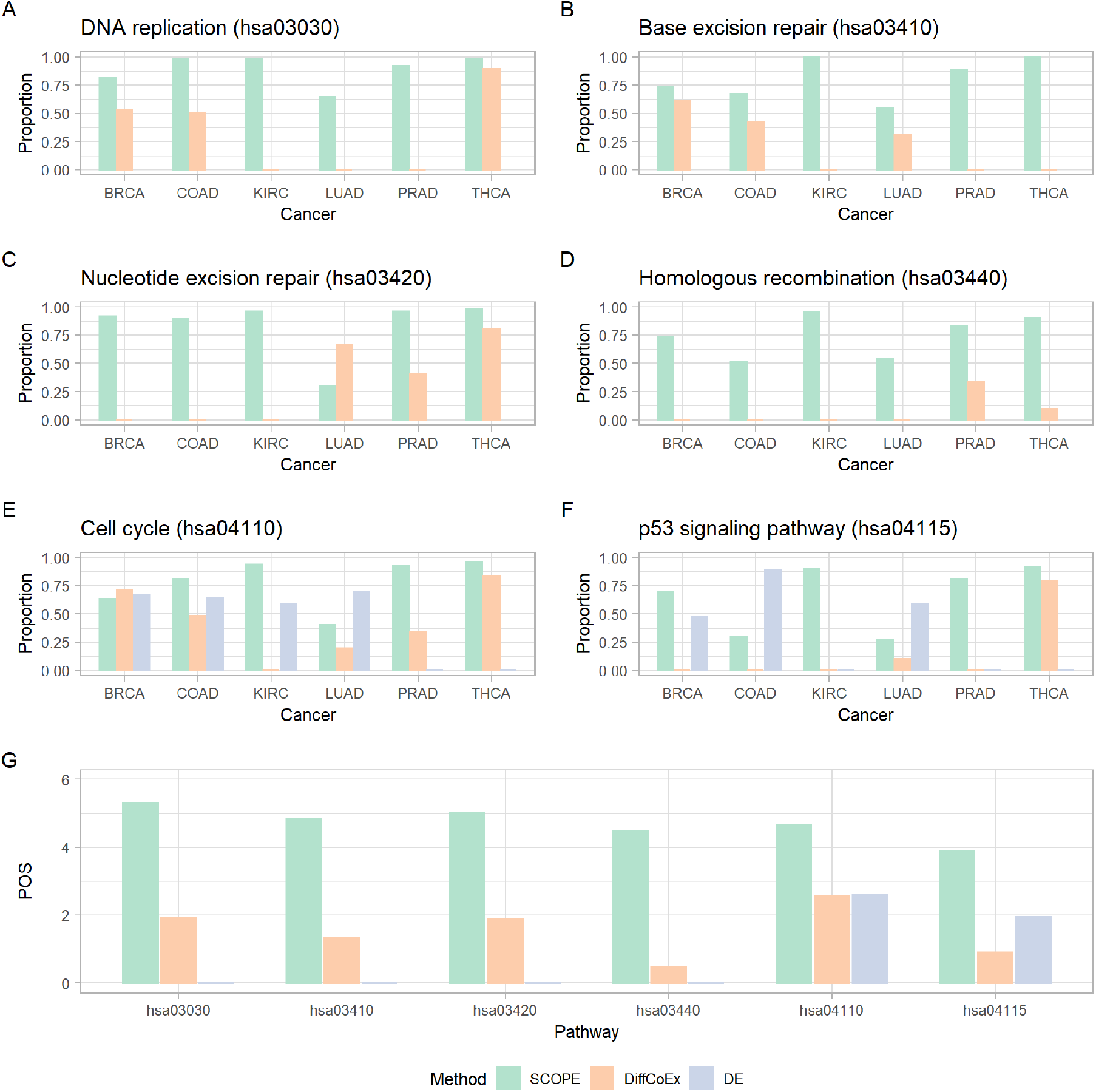
Comparison of proportion of genes in each cancer (*π*) identified by SCOPE in contrast to DiffCoEx and DE in pathways related to hallmarks of cancers. Pathways shown are **A)** DNA Replication (has03030), **B)** Base excision repair (hsa03410), **C)** Nucleotide excision repair (hsa03420), **D)** Homologous recombination (hsa03440), **E)** Cell cycle (hsa04110) and **F)** p53 signalling pathway (hsa04115). **G)** Comparison of Pathway Overlap Score (POS) across the three methods for the same pathways (**A-G**). POS is calculated as the sum of *π* across the cancers for each pathway. Higher values indicate higher discovery of genes related to each pathway across cancers.

By analyzing the literature further, we realized that the pan-cancer pathways revealed by SCOPE are biologically meaningful. Overlapped pathways enriched using Over Representation Analysis (ORA) on the KEGG Database, excluding any pathways that had no enrichment for one or more cancers, immediately reveals the high enrichment of pathways related to regulating the universal level of DNA/RNA/protein, and strikingly, the pathways related to DNA repair. The most notable characteristic of cancer is the unlimited growth of cancer cells, which also links to the Cell cycle pathway^25^. Mechanistically, they need readily available supplies of materials for cell growth and replication, e.g., more DNA replication, more RNAs transcribed by RNA polymerase and spliced by the Spliceosome, and more proteins translated by ribosome. The increased supplies are also likely due to less RNA degradation, less protein degradation by the Proteasome, and more N-glycan biosynthesis for N-linked glycosylation, one of the most abundant protein modifications which plays a critical role in tumorigenesis^26^. In addition, increased DNA replication accumulates errors as DNA mutations. Mutations inactivating tumor suppressor genes can further accelerate the accumulation of mutations, partially through defective DNA damage repair, and result in genome instability - a hallmark of all cancers^27^. Hence, this result demonstrates that the core genes identified by SCOPE-Stabilized LASSO are stably connected with pathways essential to tumor growth and/or associated with the fundamental hallmarks of any type of cancer.

### Pan-cancer pathways exhibit contrastive interaction patterns centred by core genes

To further confirm the roles of the core genes in their discovered pathways, we calculated the correlations between a core gene and all the genes in the corresponding pathway. Indeed, the core genes exhibit highly disruptive patterns in the co-expression network. Taking the nucleotide excision repair pathway network across multiple cancers as an example, the co-expression networks fundamentally differ in structure and intensity with respect to the core genes (**Figure 5A and B**). Despite different cancers employing different core genes, the correlations between the core genes and the other genes in the same pathway are universally higher or lower in the tumor tissue. More interestingly, many of the core genes are NOT differentially expressed (**Figure 5B**), indicating that core genes may contribute to cancers by disrupting their interactions although their own expression levels are not significantly altered. These results further strengthen the role that core genes appear to play in the pathology of these cancers.

**Figure 5.**
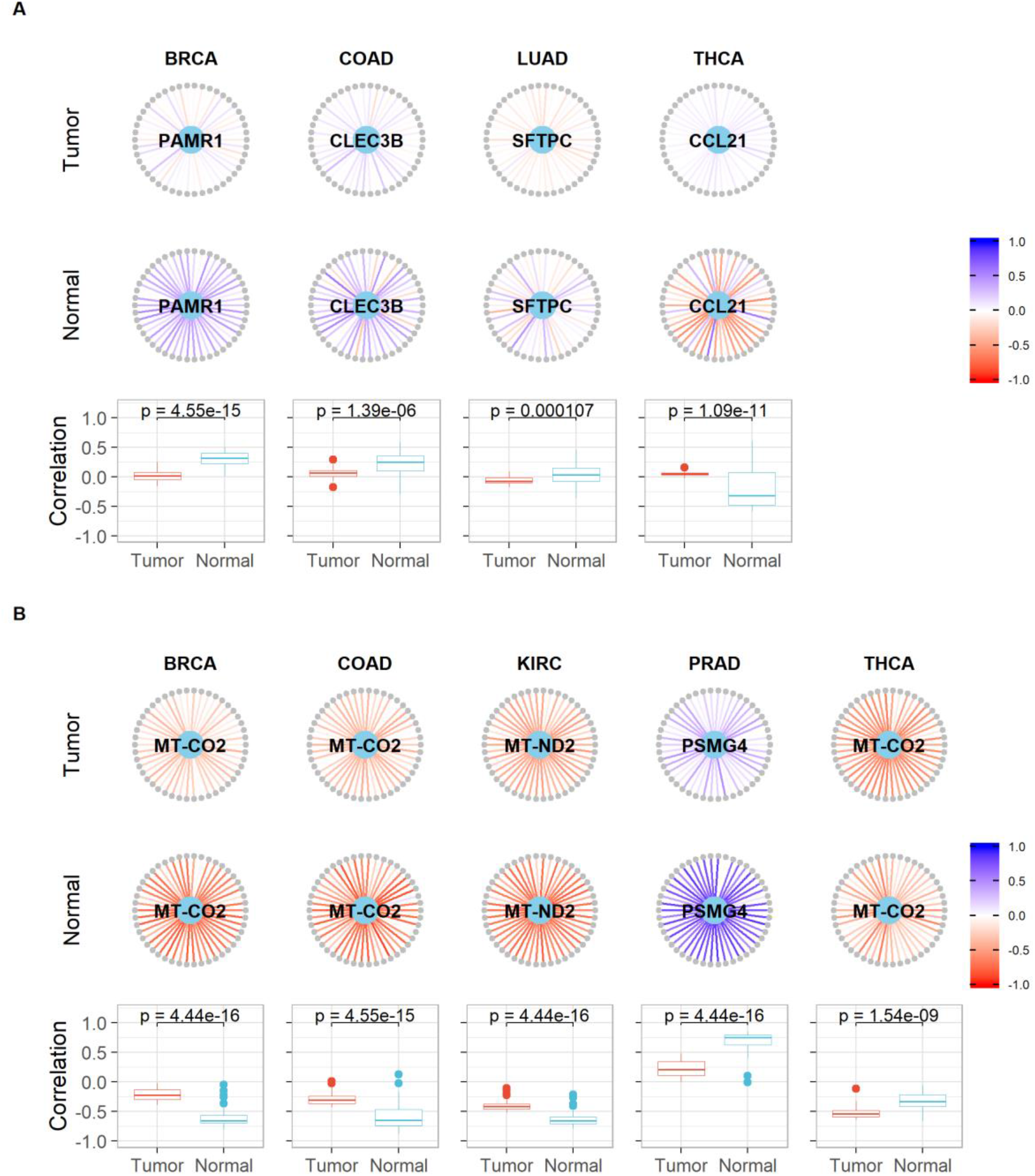
Example of roles of core genes in a pan-cancer pathway uncovered by SCOPE. The Nucleotide excision repair pathway (hsa03420) is used in this example. Core genes (light blue) are in the centre of the network with the genes in this pathway (grey) arranged in a circle. Pearson’s correlation coefficients are indicated as edges ranging from red (−1) to blue (+1). The names of the genes and their correlations with the core genes are noted in **Supplementary Table S7**. Boxplots contrast the distributions of two sets of correlations (tumor vs. normal) along with the p-value for the Kolmogorov-Smirnov test with the null hypothesis being that the two samples were drawn from the same distribution. **A)** Core genes are differentially expressed and **B)** core genes are NOT differentially expressed.

Other co-expression networks centralized by other core genes are provided in **Supplementary Figures S1-S22** and **Supplementary Tables S6**, showing switched (opposing) correlation patterns (and in some cases, an absence of correlations) when contrasting tumor tissue to normal tissue. These switched correlations appear to indicate that the core genes identified by the SCOPE-Stabilized LASSO method are highly connected genes that are indicators of the proper functioning of these pathways, if not responsible for mediating these pathways.

In addition to the above pan-cancer shared pathway analysis, SCOPE also identified cancer specific pathways, some of which show contrastive connectivity patterns. For instance, in breast cancer, Glyoxylate and dicarboxylate metabolism, Fatty acid degradation, and Regulation of lipolysis in adipocytes were discovered^28^ (**Supplementary Figure S23A, B and C**). For colon cancer, Bile secretion, Mineral absorption, and Proximal tubule bicarbonate reclamation were highlighted (**Supplementary Figure S23D, E and F**). Interestingly, out of the SCOPE identified colon cancer specific pathways, 90% are classified as being in the category of metabolism while other cancers do not show such patterns.

### In-depth annotation reveals hypothetical CD63-centred mechanism in kidney cancer

Among the five core genes selected by SCOPE in kidney cancer (KIRC), *CD63* plays an indispensable role in *VEGFR2* activation in response to *VEGF*^29^. Notably, aberrant activation of the *VEGF*-*VEGFR* axis is a pivotal driver in kidney cancer since more than 60% of kidney cancer patients harbor *VHL* mutations^30^. Inactivated *VHL* fails to degrade hypoxia-inducible factor α-subunits (*HIFα*) in kidney cancer cells. The accumulation of *HIFα* induces *VEGF* expression and secretion which further cause autocrine or paracrine activation of the *VEGFR* signaling pathway^22^. Hence *CD63* probably plays an oncogenic role in kidney cancer. Consistently, high mRNA level of *CD63* associates with adverse prognosis in KIRC patients (p=0.0019) (**Figure 6A**). In contrast, there is no such relationship in other five cancer types (**Supplementary Figure S24**). In agreement with *CD63*’s role in the activation of *VEGFR* signaling pathway, which is driven by *VHL* mutations in KIRC, the association is more significant in *VHL*-mutated cohorts (p=0.0006) (**Figure 6B**), than in *VHL*-wildtype cohorts (p=0.216) (**Figure 6C**). Along this line, high expression of *CD63* in kidney tumors correlates with a hypoxia gene signature assessed by two different scores (**Figure 6D and E**). *CD63* is also known as a marker of exosomes, extracellular vesicles secreted by cells^31^. In agreement with the fact that exosomes can contribute to metastasis^32^, *CD63* shows a tendency to be correlated with metastasis in KIRC patients although barely above the significant cut-off of 0.05 (p=0.0565) (**Figure 6F**). In particular, *CD63* knockout mice are viable, fertile and almost normal except for an altered water balance, such as increased urinary flow, water intake, reduced urine osmolality, and a higher fecal water content^33^. This does not only suggest that *CD63* plays a critical and specific role in kidney pathology, and consequently in kidney tumorigenesis, but also hints that *CD63* can be a therapeutic target for kidney cancer with minimal systemic toxicity. Anti-*CD63* antibodies were reported to suppress allergy^34^ or inhibit metastasis^35^ *in vivo*. It will be worthy exploring whether anti-*CD63* antibodies are able to improve the potency of targeted therapy or immunotherapy and inhibit metastasis in kidney cancer patients. Nevertheless, this example suggests the core genes selected by SCOPE may help exert bona fide biological functions in the mechanisms of cancer.

**Figure 6.**
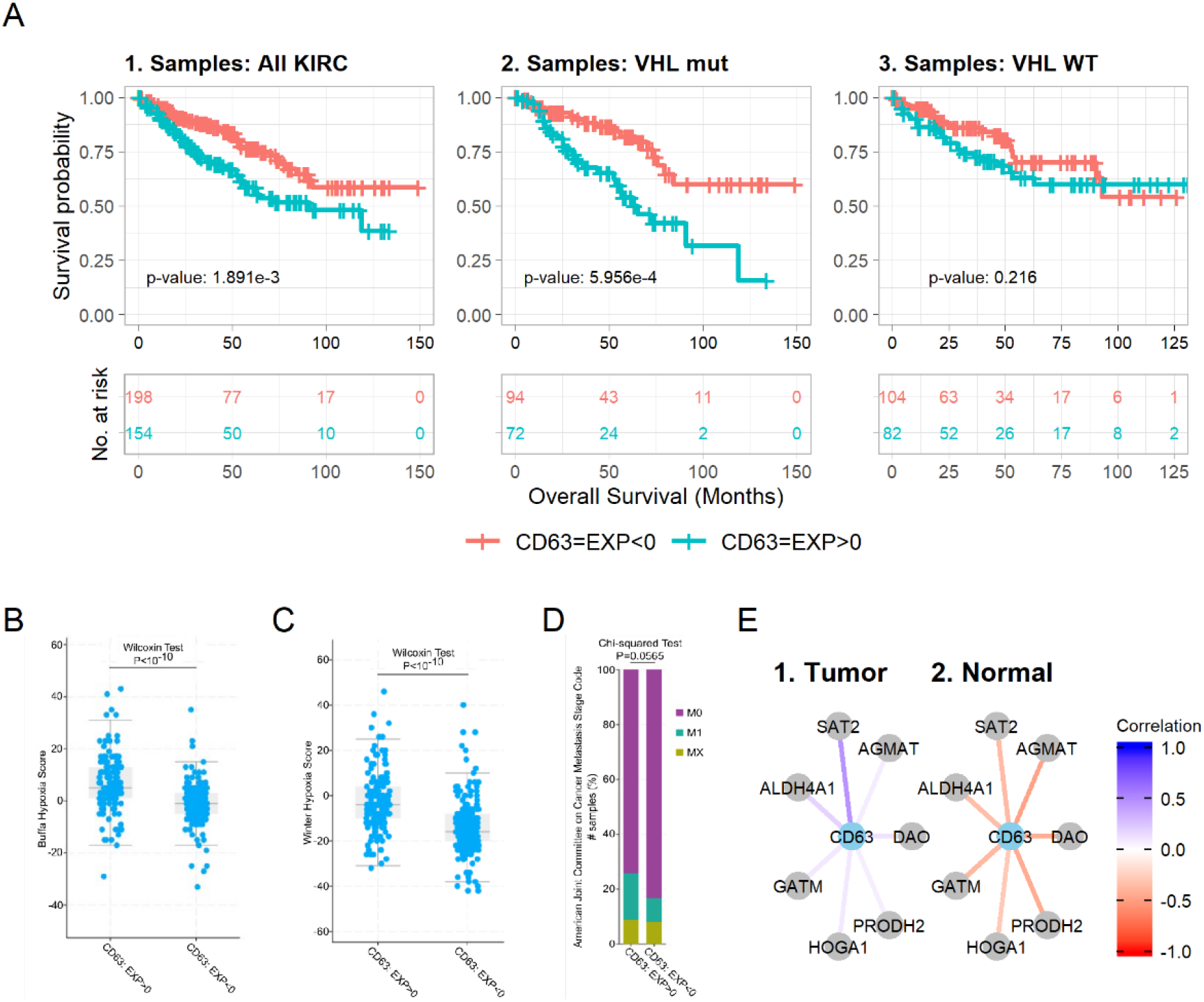
Hypothetical role of *CD63* in kidney cancer. EXP < 0 indicates samples in which the expression level of the gene (*CD63*) is lower than the arithmetic mean of the expression levels of the gene across all samples; while EXP > 0 indicates higher than mean expression levels. Survival plots of patients considering differing expression of *CD63* in **A.1)** all patients in the KIRC dataset, **A.2)** patients with *VHL* mutation and **A.3)** patients with *VHL* wildtype. **B)** and **C)** indicate that a higher expression of *CD63* correlates with well known hypoxia gene signatures while **D)** indicates the relationship between *CD63* and metastasis in kidney cancers. **E.1)** and **E.2)** connectivity network suggesting the role *CD63* plays in the arginine and proline metabolism pathway with key genes involved in the pathway. (Data and figures of Panels **A - E** are derived from the cBioPortal website (https://www.cbioportal.org/).

Among all genes connected with *CD63, SAT2* is the one with the most significantly differential correlations in tumor and normal tissues. *SAT2* mRNA level shows a negative correlation with CD63 in normal tissue while exhibiting an almost opposite correlation in tumors (**Figure 6G and H**). Many other genes in the pathway of arginine and proline metabolism, such as *AGMAT, DAO, ALDH4A1, PRODH2, GATM* and *HOGA1*, also show similar patterns of switched correlations with *CD63* (**Figure 6G and H**). The altered correlations in tumors uncovered by SCOPE hint that these genes may play critical roles in kidney tumorigenesis. In agreement with this hypothesis, agmatinase, encoded by *AGMAT*, is diminished in kidney cancer samples whereas *AGMAT* mRNA is most abundant in human liver and kidney^36^. Moreover, *SAT2, DAO, ALDH4A1, PRODH2, GATM* and *HOGA1* are ubiquitously expressed in kidney based on the Human Protein Atlas^37,38^. However, other genes belonging to the pathway of arginine and proline metabolism were not identified, or only shown negligible correlation differences by SCOPE, such as *SAT1, NOS1, CKM, CKB* and *ARG2*, and do not show obvious overexpression in kidney^37–39^. The distinct tissue specificity of two groups of genes in the same pathway of arginine and proline metabolism validates that SCOPE was able to identify altered co-expression patterns in specific cancer types. In contrast, neither DE nor DiffCoEx uncovered significant enrichment of these pathways in KIRC, further strengthening the ability of SCOPE in uncovering such biologically relevant pathways.

## Discussion

Current methods of driver gene identification utilize multi-omics data, in particular mutation data in collaboration with known biological pathways. Transcriptomic data are seldom used for identification of driver genes. This is in part due to the inability to determine causality using methods such as differential expression, differential co-expression and co-expression networks. Gene expression data alone, while conveniently available, are infrequently used for this purpose and rather directed towards biomarker discovery. Our proposed method of stabilizing the LASSO such that it identifies consistent predictors followed by co-expression and pathway analysis enables researchers to identify the core genes and pathways by taking advantage of the synergy between two disconnected fields: linear feature selection and nonlinear co-expression network analysis. This provides a method for experimentalists to narrow down candidate genes using more cost-effective expression data. Furthermore, since only a handful of such core genes are selected – this enables experimentalists an ideal scenario of being able to study these few genes extensively.

We note that using a SCOPE-Stabilized LASSO with a stringent threshold not only allows for the selection of stable predictors – but also a smaller number of genes that could be functionally validated experimentally. The choice of this threshold is not too different from the requirement of co-expression networks and differential co-expression networks to choose the *β* power parameter (higher values emphasise larger differences in correlation) – a more stringent threshold will reduce predictive power but increase the likelihood of retaining genes that potentially are drivers of the disease. In the six cancers analyzed here, we note that breast invasive carcinoma, in particular, has a large number of identifiable core genes– stable predictors across samples. Thus, a more stringent threshold of *θ*_*thr*_ = 0.90 was assigned to this dataset while the others had *θ*_*thr*_ = 0.75. These selections are validated in **Supplementary Table S2** where we see that these selections are stable even across multiple SCOPE-Stabilized LASSOs.

Discovering enriched pathways connected with core genes may provide rationale for targeted therapy against certain cancer types. A series of genes in the ferroptosis pathway, including *ACSL4, MAP1LC3B, ATG5, PRNP, NCOA4, PCBP1, LPCAT3, VDAC3, FTH1, SLC39A14, SLC40A1* and *SLC11A2*, showed significantly changed patterns of correlation with PSMG4 in the PRAD dataset (**Supplementary Table S7**). Ferroptosis is a programmed cell death driven by iron-dependent phospholipid peroxidation and ROS generation^40^. Since excessive iron contributes to ferroptosis, *PCBP1* and *FTH1*, which regulate iron metabolism and storage, are considered negative regulators of ferroptosis. *ATG5, MAP1LC3B* and *NCOA4* initiate autophagy and consequently promote iron-release from degraded iron-bound proteins. *SLC40A1, SLC39A14* and *PRNP* export iron from cells and reduce ferroptosis whereas *SLC11A2* regulates iron release to the cytoplasm and may enhance ferroptosis. *ACSL4, LPCAT3* and *VDAC3* regulate the mechanism of phospholipid and *NADH* oxidation and play roles as positive regulators of ferroptosis^41^. In particular, *AIFM2*, a critical ferroptosis suppressor identified in 2019^42,43^, shows reduced expression in prostate cancer (PRAD) (logFC = -0.9008). All these data indicate that ferroptosis inducers might be potent in PRAD patients. Consistently, recent work has reported induction of ferroptosis as a new therapeutic strategy for advanced prostate cancer^44^. Neither DiffCoEx nor DE highlighted the ferroptosis pathway as significant in PRAD while SCOPE was able to highlight this pathway uniquely and significantly in PRAD.

An inherent limitation of transcriptomic data is that most biological functions are performed by proteins, not mRNAs. One example is the p53 signaling pathway in BRCA, which is significantly enriched by the gene pairs of *MT-CO2* with *CDK4, AIFM2* or *CHEK2* (**Supplementary Table S8**). Furthermore, another core gene, *CD300LG*, shows switched correlations with *TP53* and *CASP9* even though the respective CGN is not enriched for the p53 signaling pathway. Although *TP53* (encoding p53) showed altered correlations with *CD300LG* and *MT-CO2*, the putative transcriptional targets of p53, such as *CDKN1A* and *MDM2*^45^, did not show significant changes of correlation. It implies the p53 transcriptional activity was not significantly changed in the presence of significantly changed mRNA level of *TP53*. This implication was further supported by two facts: (1) the regulation of p53 activity is dominant at the post-translational level47474747, not the mRNA level; (2) 35% of patients in the TCGA-BRCA database harbor *TP53* mutations and most *TP53* mutations abolish the transcriptional activity of p53. We looked for top transcription factor binding sites in the promoters of these genes (*CASP9, CDK4, AIFM2* and *CHEK2*) provided by QIAGEN through Genecards^46^, and found CCAAT-enhancer-binding proteins (C/EBPs) bound to these promoters. Since the *PI3K*-*AKT*-*mTOR* signaling pathway is highly mutated in the BRCA database (**Supplementary Figure S25**, obtained from cBioPortal in the TCGA-BRCA database^47,48^) and is able to regulate the transcriptional activity of C/EBPs^49^, a reasonable explanation is that a hyperactivated *PI3K*-*AKT*-*mTOR* axis induces the mRNA expression of these targets via C/EBPs as the transcriptional factor (but not *TP53*) in BRCA patients. Nevertheless, with more data of cancer at protein level, such as The Pathology Atlas^38^ and The Cancer Proteome Atlas Portal^50,51^, SCOPE may be substantially empowered to provide more valuable insights on the aberrant connections in tumor cells.

To recap, we have presented SCOPE, a method stabilizing gene selection and co-expression network analysis, which is able to identify core genes and pathways underlying cancers. Its effectiveness has been demonstrated by various analysis from three angles (i.e., selection of few, stable and predictive genes, pan-cancer shared pathways, and the role of core genes in connectivity analysis). Moreover, in-depth annotations have revealed the pivotal role of *CD63* on tumorigenesis in kidney cancer and potential therapeutic application of anti-*CD63* antibody on kidney cancer patients. As a proof of concept, we have only contrasted cancer and normal tissues in this work. However, the statistical framework is applicable to any case/control settings. In the future, we will adapt SCOPE to analyze clinically important qualities such as whether a patient will respond to medical treatments such as immunotherapy, paving the way to the application of precision medicine in more applications.

## Acknowledgement

This work is partly supported by NSERC Discovery grant (RGPIN-2018-05147), New Frontiers in Research Fund (NFRFE-2018-00748) and VPR Catalyst grant to Q.Z., and Alberta Cancer Foundation to C.S.S. (Grant # 27246). The computational infrastructure to conduct the analysis is supported by a Canada Foundation for Innovation JELF grant (36605). The authors are grateful to Dr. Jurg Ott and Dr. Xingyi Guo for their comments.

## Methods

### SCOPE-Stabilized LASSO Selection

While the LASSO has proven versatile in many applications, statistically it has become apparent that in the presence of multiple correlated features it may be inconsistent in its selection of features even in multiple random samplings of the same data^52^. This has led to a number of new methods being proposed that are all modifications of the original LASSO such as adaptive LASSO^52^, random LASSO^53^, bolasso^19^, etc. A seminal work by Meinshausen and Bühlmann (2010)^17^ discusses stability paths – which obtain the selection probabilities of each feature by subsampling along all possible values of the tuning parameter with randomized LASSO which introduces a random penalty *λ* for each feature. While such methods are statistically proven and can lead to sound results, they appear to be seldom used in the field of genomics.

In SCOPE, a simpler solution to the inconsistency of variable selection in LASSO is proposed and applied in the form of a bootstrapped LASSO [**Figure 1A and B**]. While simple in design, it produces consistent results which are highly predictive. In the case of this paper, where the phenotype is binary (tumour or normal sample), a logistic LASSO regression model is trained multiple times by subsampling from the same dataset. Genes that were selected in most of the models (over a threshold proportion *θ*_*thr*_) are used to build a final logistic regression model for which the final accuracy will be assessed. This stable subset of genes is proposed to be the “core” genes of the disease. These core genes can then be used in other predictive models or for further downstream analysis; SCOPE employs a co-expression-based pathway analysis utilizing these selected core genes.

The SCOPE-Stabilized LASSO used in this analysis features the consensus of 1,000 training-test splits of a 70%-30% split ratio (each with a consistent case/control ratio of the full dataset). Each LASSO model trained was tuned for the optimal value of *λ* using 10-fold cross-validation. The thresholds used for the different datasets of the real analysis are detailed in **Results**.

### Co-expression and Pathway Analysis

It is assumed that core genes interact with multiple other genes which may be involved in pathways responsible for disease mechanisms. To identify these genes, we conducted a co-expression analysis (**Figure 1D**). To claim a gene as being significantly co-expressed with a core gene, we required a null distribution for the correlations (co-expression) between pairs of random genes. To this end, we drew random pairs of genes and calculated the Pearson Correlation Coefficients of these pairs. Using this distribution, we obtain the 97.5^th^ percentiles for both positive and negative correlations. This allowed us to identify genes that are significantly co-expressed with core genes. Each set of genes thus identified (secondary co-expressed genes, along with their corresponding core gene) termed Core Gene Networks (CGNs) were then tested for pathway enrichment.

To reflect the fact that some critical genes are not so highly co-expressed, are however, significantly differentially co-expressed when contrasting cancer and normal tissues, we also obtained the genes that are significantly differently co-expressed with the core gene between tumor and normal tissues (**Figure 1D**). As in the case of the co-expression analysis, a null distribution of the differential co-expression values (|*corr*_*case*_ − *corr*_*control*_|) was obtained and the 97.5^th^ percentile was used to select significantly differentially co-expressed secondary genes. These secondary genes from differential co-expression analysis were also added to the CGNs for pathway analysis below.

Pathway enrichment is typically used to assess whether a particular set of genes overlap with known biological pathways significantly higher than by chance. There are many databases containing such pathways, and SCOPE employs the Kyoto Encyclopedia of Genes and Genomes (KEGG)^54,55^ database due to its comprehensiveness and popularity.

Over Representation Analysis^56^ is used to identify the statistical significance and the R package, *WebGestaltR*^57,58^ was used for testing pathway enrichment against the KEGG database. This analysis results in several pathways enriched (at a False Discover Rate (FDR) ≤ 0.05) for each CGN underlying the focal core gene. SCOPE then discovers pathways that are commonly influenced by CGNs (seeded by different core genes).

For a single disease such as a cancer, an index for the level of sharing of a pathway (across multiple CGNs within a cancer), *θ*_*cancer*_, is defined as the number of co-expressed genes (including the core gene) found to be enriched in this pathway divided by the total number of the genes in the pathway. When multiple diseases are jointly analyzed (e.g., the six cancers used in this paper as a demonstrating example), SCOPE will further discover pathways common to all diseases (**Supplementary Table S5**). The summation of this single cancer-specific index (*θ*_*cancer*_) over all the cancers is noted as the Pathway Overlap Score (POS). Intuitively, a higher POS score indicates a higher overlap of the pathway across different cancers.

### Methods compared to SCOPE

#### LASSO (Least Absolute Shrinkage and Selection Operator)

The primary benchmark and point of comparison is a traditional *L*_*1*_ regularized logistic regression model which uses the addition of the absolute value of the coefficients to promote sparsity in the loss function. Given *n* number of samples and *p* number of features/variables, the regularized loss function of a logistic regression model takes the form:

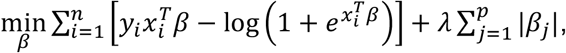

where *λ* is the tuning parameter controlling the trade-off between sparsity and accuracy. 10-fold cross-validation is typically employed to tune for this parameter and the R package *glmnet*^59,60^ enhanced by *glmnetUtils*^61^ was used to fit LASSO models in this paper (**Supplementary Materials**). Genes selected by a traditional LASSO to genes selected by SCOPE-Stabilized LASSO step are compared in the **Results** section.

#### Differential Co-expression Analysis (DiffCoEx)

Standard network-based methods of analysis of expression data typically utilize interconnectedness of genes (in the form of co-expression) to identify important networks (or “modules”) of genes. However, in a case-control setting, the use of differential co-expression can prove more informative due to the contrastive nature of the analysis. DiffCoEx^62^, one of the most popular methods extending the popular WGCNA co-expression network analysis tool was chosen for comparison. DiffCoEx identified differentially co-expressed modules (**Supplementary Materials**) which were used in pathway enrichment similar to how CGNs were used for pathway enrichment and studying pathway overlaps.

#### Differential Expression (DE)

An edgeR-limma based pipeline^63^ was used to normalize the data to log2-counts per million (log-CPM) values and a linear model incorporating weights from *voom* to correct for the mean-variance relationship was used to statistically detect differential expression of genes in each of the cancers. The pipeline was run using default values for all parameters as described in the workflow.

### Real Data Analysis

#### Data Source and Processing

The Cancer Genome Atlas (TCGA) Program initiated by the National Cancer Institute and the National Human Genome Research Institute in 2006^20^ offers a wealth of omics data on 32 different cancers and their subtypes at 68 primary sites. This includes RNA-Seq data which provides a snapshot of the transcriptomic landscape of the tumor site as well as of solid normal tissue close to the tumor site.

Six cancers (breast invasive carcinoma – BRCA, kidney renal clear cell carcinoma – KIRC, lung adenocarcinoma – LUAD, colon adenocarcinoma – COAD, prostate adenocarcinoma – PRAD and thyroid carcinoma - THCA) were chosen primarily due to the large samples of expression data available as well as the inclusion of normal tissue from the same patients. A breakdown of the sample sizes and disease status is given in **Supplementary Table S9**.

The raw count data was downloaded from the TCGA data portal and then converted to Transcripts Per Million (TPM) values using gene lengths obtained through the *BioMart*^64,65^ package. Phenotype (tumor or normal) was determined based on the Sample Type column provided in the database and “Primary Tissue” was considered as cases and “Normal Tissue” as controls. Any other sample types such as “Metastasis” were discarded. Models were then fitted on this processed data.

Two additional datasets (lung cancer associated), E-MTAB-6043^23^ and MTAB-6699^24^ were downloaded from ArrayExpress and used to externally validate the predictive accuracy of core genes selected by SCOPE and alternative methods.

