## Supplementary Materials for "Stabilized COre gene and Pathway Election uncovers pan-cancer shared pathways and a cancer specific driver"

**SCOPE Algorithm (Extended)**

**Input:** Expression Data $\mathbf{X}_{n\times p}$
Phenotype Data $\mathbf{Y}_{n\times 1}$
Number of splits $m$
Proportion of selection threshold $\theta_{thr}$

Stabilized LASSO
**for** $i = 1$ to $m$
  Generate a random train-test split
  Calculate LASSO estimates $\beta_{ji}$ for split $i$
end for
Compute $\theta_{\beta_j} = \frac{\left|\sum_{\mathrm{i}=1}^{m} I(\beta_{ji} \neq 0)\right|}{m}$, where I(.) is an indicator function
Select core genes $CGs = \left\{gene_j \middle| \theta_{\beta_j} \geq \theta_{thr}\right\}$

Co-expression Analysis
**for** $i = 1$ to $1000$
  Select 1000 random genes
  Calculate correlation values
  $\boldsymbol{corrs_{thr}^{+}}, \boldsymbol{corrs_{thr}^{-}}.\mathrm{append}$(97.5th percentile of positive and negative correlations)
**end for**
$corr_{thr}^{+}, corr_{thr}^{-} = \mathrm{medians}(corrs_{thr}^{+}, corrs_{thr}^{-})$
**for each** $\boldsymbol{CG_i}$
  Calculate $corr_{i,j}$ ∀ other genes $j$
  **if** $corr_{i,j} > corr_{thr}^{+}$ $\mathrm{OR}$ $corr_{i,j} < corr_{thr}^{-}$ **then**
    $gene_j$ is co-expressed with core gene $i$
    $\mathrm{CGN}_i.\mathrm{append}(gene_j)$
end for
Repeat for Differential Co-expression

Pathway Enrichment
**Input:** $\mathrm{CGN}_i\ \forall i$
for each $\mathbf{CGN}_{\boldsymbol{i}}$
  $P_i = \{p | p \text{ enriched in } \mathrm{CGN}_i\}$
**end for**
  $\forall\, p_j, PS_j = \sum_{i=1}^{|CG|} I(p_j \in P_i)$

Output:
Pathways enriched for each $\mathrm{CGN}_i, \boldsymbol{P_i}$
Number of CGNs enriching each pathway $p_j, \boldsymbol{PS_j}$

### LASSO (Least Absolute Shrinkage and Selection Operator)

Among multiple methods proposed for variable selection and model building, the LASSO[1] and ridge[2] regularized regression methods have gained popularity in recent years for high-dimensionality problems. Regularization typically refers to the addition of an additional term in the loss function that is meant to prevent overfitting. LASSO (a form of $L_1$ regularization) adds the sum of absolute values of the regression coefficients to the loss function and in turn allows some of these coefficients to be reduced to 0 – thus inducing sparsity. It has been shown that L1 regularization promotes sparsity more than higher order regularizations (such as L2, or ridge regression) among convex forms of regularization models[1].

In particular, a typical regularized loss function is:

$$\min_{\beta} \sum_{i=1}^{n} \left(y_i - x_{\mathrm{i}}^T \beta\right)^2 + \lambda \sum_{j=1}^{p} |\beta_j| \,,$$

where $\lambda$ is a non-negative tuning parameter that controls the trade-off between sparsity and accuracy, $n$ is the number of samples and $p$ is the number of features/variables.

In this paper, a logistic regression model is used and therefore, the loss function becomes,

$$\min_{\beta} \sum_{i=1}^{n} \left[y_i x_i^T \beta - \log\left(1 + e^{x_i^T \beta}\right)\right] + \lambda \sum_{j=1}^{p} |\beta_j|.$$

The tuning parameter $\lambda$ is typically chosen via cross-validation and in this paper, a 10-fold cross validation is used to select the optimal $\lambda$.

### Differential Co-expression Analysis (DiffCoEx)

To compare our method with a standard method based on network analysis, we use differential co-expression networks to identify groups (or “modules”) of differentially co-expressed genes and conduct pathway enrichment on these modules. While there are multiple methods of differential co-expression analysis, the most widely used method remains DiffCoEx[3], an extension of the popular WGCNA[4] to differential co-expression. DiffCoEx begins with the construction of two adjacency matrices: $C_{case}: c_{ij}^{case} = cor(gene_i, gene_j)$ for cancerous samples and $C_{control}$ similarly for healthy samples.

While different correlation measures can be used in this step, the authors of DiffCoEx used the Spearman rank correlation. A matrix of adjacency difference is then calculated,

$$D: d_{ij} = \left(\sqrt{\frac{1}{2}\left|sign\left(c_{ij}^{case}\right) * \left(c_{ij}^{case}\right)^2 - sign\left(c_{ij}^{control}\right) * \left(c_{ij}^{control}\right)^2\right|}\right)^{\beta}$$

where $\beta \geq 0$ is an integer tuning parameter which can be selected in multiple ways. In this paper we choose $\beta \in [5,6,7,8,9,10]$ such that each cancer has the minimum number of modules with the largest module containing the smallest number of genes. Next, a Topological Overlap dissimilarity Matrix (TOM) is calculated where lower values of $t_{ij}$ indicate that a pair of genes $gene_i$ and $gene_j$ have significant correlation changes (between case and control) with the same group of genes.

$$T: t_{ij} = 1 - \left(\frac{\sum_k d_{ik} d_{kj} + d_{ij}}{\min\left(\sum_k d_{ik}, \sum_k d_{jk}\right) + 1 - d_{ij}}\right)$$

Finally, the dissimilarity matrix $T$ is used for clustering and “modules” of differentially co-expressed genes are identified. These modules which contain sets of genes are then tested for pathway enrichment and the same measures of overlap as in the SCOPE method are calculated.

### Supplementary Tables

**Supplementary Table S1 – Core genes identified for each of the 6 selected cancers of the TCGA database.** The $\theta$ value of each gene denotes the proportion of models in which the gene was selected. Since this was based on 1,000 independent LASSO runs, i.e., a value of 0.779 for GYPE gene in LUAD indicates that GYPE was selected by 779 of the LASSO models run for the LUAD dataset. Core genes selected based on the $\theta_{thr}$s are indicated in bold.

| BRCA | | COAD | | KIRC | | LUAD | | PRAD | | THCA | |
|---|---|---|---|---|---|---|---|---|---|---|---|
| Gene | $\theta_{BRCA}$ | Gene | $\theta_{\mathbf{COAD}}$ | Gene | $\theta_{\mathbf{KIRC}}$ | Gene | $\theta_{\mathbf{LUAD}}$ | Gene | $\theta_{\mathbf{PRAD}}$ | Gene | $\theta_{\mathbf{THCA}}$ |
| **HNRNPL** | 1.000 | **SCARA5** | 1.000 | **FAM245A** | 0.973 | **EXOSC1** | 0.999 | **MT-CO2** | 1.000 | **RPS27L** | 1.000 |
| **PAMR1** | 1.000 | **TRIM27** | 1.000 | **MT-ND2** | 0.958 | **SFTPC** | 0.998 | **APOBEC3C** | 1.000 | **MT-CO2** | 1.000 |
| **VEGFD** | 1.000 | **ANAPC7** | 0.999 | **CORO7** | 0.913 | **GMPPA** | 0.938 | **ALDH1L1-AS2** | 0.828 | **CTSA** | 0.984 |
| **OST4** | 1.000 | **IRGQ** | 0.873 | **CD63** | 0.810 | **PMM2** | 0.932 | **CA14** | 0.773 | **CCL21** | 0.965 |
| **MT-CO2** | 0.996 | **TMIGD1** | 0.871 | **STX4** | 0.753 | **STX11** | 0.874 | **PSMG4** | 0.755 | **LINC01834** | 0.879 |
| **TSLP** | 0.993 | **CLEC3B** | 0.814 | MPV17 | 0.736 | **GPD1** | 0.785 | AAAS | 0.734 | **DPT** | 0.876 |
| **CD300LG** | 0.991 | **MT-CO2** | 0.791 | ANKRD39 | 0.726 | **GYPE** | 0.779 | GPR89B | 0.719 | **AL034374.1** | 0.763 |
| **OXTR** | 0.988 | BMP3 | 0.747 | TCF21 | 0.680 | **FABP4** | 0.771 | POLR2H | 0.601 | **MMRN1** | 0.761 |
| **ARF1** | 0.961 | CA2 | 0.716 | GGT6 | 0.664 | LINC01996 | 0.743 | GAR1 | 0.599 | **RYR2** | 0.756 |
| **RNF149** | 0.955 | MT-ND4 | 0.632 | DTX2 | 0.644 | C1orf131 | 0.688 | RAB17 | 0.536 | RELN | 0.683 |

**Supplementary Table S2 – Stability of the SCOPE-Stabilized LASSO step.** Among 100 runs, the proportion of runs selecting the gene as a core gene is listed in the right half of each column. "Prop" value of each gene denotes the proportion of SCOPE-Stabilized LASSO models that the gene was selected in. Since 100 SCOPE-Stabilized LASSO runs were used to obtain the results shown in this table, i.e., a value of 0.83 for FABP4 in LUAD indicates this gene was selected in 83 of the SCOPE-Stabilized LASSO runs for LUAD. Underlined genes are genes that were not selected in the SCOPE-Stabilized LASSO run used for the analysis in this paper.

| **BRCA** | | **COAD** | | **KIRC** | | **LUAD** | | **PRAD** | | **THCA** | |
|---|---|---|---|---|---|---|---|---|---|---|---|
| **Gene** | **Prop** | **Gene** | **Prop** | **Gene** | **Prop** | **Gene** | **Prop** | **Gene** | **Prop** | **Gene** | **Prop** |
| HNRNPL | 1.00 | IRGQ | 1.00 | CD63 | 1.00 | STX11 | 1.00 | CA14 | 1.00 | CTSA | 1.00 |
| ARF1 | 1.00 | SCARA5 | 1.00 | MT-ND2 | 1.00 | PMM2 | 1.00 | MT-CO2 | 1.00 | CCL21 | 1.00 |
| TSLP | 1.00 | TMIGD1 | 1.00 | FAM245A | 1.00 | GMPPA | 1.00 | APOBEC3C | 1.00 | DPT | 1.00 |
| PAMR1 | 1.00 | ANAPC7 | 1.00 | CORO7 | 1.00 | GPD1 | 1.00 | ALDH1L1-AS2 | 1.00 | RPS27L | 1.00 |
| CD300LG | 1.00 | MT-CO2 | 1.00 | STX4 | 0.69 | SFTPC | 1.00 | AAAS | 0.53 | MT-CO2 | 1.00 |
| RNF149 | 1.00 | TRIM27 | 1.00 | ANKRD39 | 0.48 | EXOSC1 | 1.00 | PSMG4 | 0.37 | LINC01834 | 1.00 |
| VEGFD | 1.00 | CLEC3B | 0.99 | MPV17 | 0.30 | GYPE | 1.00 | GPR89B | 0.01 | MMRN1 | 0.90 |
| OXTR | 1.00 | CA2 | 0.54 | | | LINC01996 | 0.87 | | | AL034374.1 | 0.90 |
| MT-CO2 | 1.00 | BMP3 | 0.23 | | | FABP4 | 0.83 | | | RYR2 | 0.37 |
| OST4 | 1.00 | | | | | | | | | | |

**Supplementary Table S3 – Within cancer pathway overlap across Core Gene Networks.** The number of CGNs that uncovered each pathway as enriched in each cancer is given in the table. This table shows the top 10 pathways sorted by the number of CGNs the pathway was uncovered in.

| KEGG GeneSet | Pathway Name | BRCA ( /10) | COAD ( /7) | KIRC ( /5) | LUAD ( /8) | PRAD ( /5) | THCA ( /9) | Total ( /44) |
|---|---|---|---|---|---|---|---|---|
| hsa04110 | Cell cycle | 5 | 4 | 2 | 5 | 4 | 2 | 22 |
| hsa03460 | Fanconi anemia pathway | 5 | 4 | 1 | 5 | 2 | 1 | 18 |
| hsa00190 | Oxidative phosphorylation | 6 | 2 | 4 | 2 | 1 | 2 | 17 |
| hsa03030 | DNA replication | 5 | 4 | 2 | 4 | 1 | 1 | 17 |
| hsa05012 | Parkinson disease | 6 | 2 | 3 | 2 | 1 | 3 | 17 |
| hsa00240 | Pyrimidine metabolism | 5 | 4 | 2 | 3 | 1 | 1 | 16 |
| hsa00510 | N-Glycan biosynthesis | 4 | 2 | 1 | 5 | 1 | 3 | 16 |
| hsa03050 | Proteasome | 6 | 4 | 1 | 2 | 1 | 2 | 16 |
| hsa03410 | Base excision repair | 5 | 3 | 1 | 4 | 1 | 2 | 16 |
| hsa04141 | Protein processing in endoplasmic reticulum | 5 | 1 | 1 | 4 | 2 | 3 | 16 |
| hsa04714 | Thermogenesis | 6 | 2 | 2 | 2 | 2 | 2 | 16 |
| hsa03040 | Spliceosome | 5 | 4 | 1 | 3 | 1 | 1 | 15 |
| hsa03440 | Homologous recombination | 5 | 3 | 2 | 3 | 1 | 1 | 15 |
| hsa03420 | Nucleotide excision repair | 4 | 4 | 1 | 1 | 2 | 2 | 14 |
| hsa04115 | p53 signaling pathway | 1 | 3 | 2 | 1 | 1 | 6 | 14 |
| hsa03013 | RNA transport | 4 | 4 | 1 | 2 | 1 | 1 | 13 |

**Supplementary Table S4 – Within cancer pathway overlaps across modules using DiffCoEx.** The number of modules that uncovered each pathway as enriched in each cancer is given in the table. This table shows the top 10 pathways sorted by the number of modules the pathway was uncovered in.

| Pathway Name | BRCA ( /3) | COAD (/17) | KIRC ( /10) | LUAD (/52) | PRAD ( /7) | THCA ( /33) | Total ( /122) |
|---|---|---|---|---|---|---|---|
| Cytokine-cytokine receptor interaction | 1 | 1 | 0 | 3 | 1 | 3 | 9 |
| AGE-RAGE signaling pathway in diabetic complications | 1 | 1 | 1 | 3 | 1 | 1 | 8 |
| Cell cycle | 1 | 2 | 0 | 2 | 1 | 2 | 8 |
| Epstein-Barr virus infection | 0 | 2 | 0 | 3 | 1 | 2 | 8 |
| Human T-cell leukemia virus 1 infection | 0 | 1 | 1 | 3 | 1 | 2 | 8 |
| Osteoclast differentiation | 1 | 1 | 1 | 2 | 1 | 2 | 8 |
| Spliceosome | 1 | 1 | 0 | 2 | 2 | 2 | 8 |
| Autophagy | 0 | 2 | 0 | 2 | 1 | 2 | 7 |
| Cellular senescence | 1 | 1 | 1 | 2 | 1 | 1 | 7 |
| Chagas disease (American trypanosomiasis) | 1 | 1 | 0 | 3 | 1 | 1 | 7 |

**Supplementary Table S5 – Distribution of proportion of genes identified by each SCOPE, DiffCoEx and DE across all pathways in each cancer along with their Pathway Overlap Scores (POS).** $\pi_{cancer}$ values denote the proportion of genes contained in CGNs (for SCOPE), contained in differentially co-expressed modules (for DiffCoEx) and contained in the list of differentially expressed genes (for DE) out of the total number of genes in each pathway the proportion of genes. POS score is the summation of this value over all cancers for each method. Higher values indicate higher overlap of pathway across cancers.
(Refer Supplementary Excel file)

**Supplementary Table S6 – Table of core genes and secondary genes involved in each of the pathways shown in Figures S1-S22**. Pearson Correlation Coefficients of gene pairs are included for each cancer and tissue type.
(Refer Supplementary Excel File)

**Supplementary Table S7 – Correlation patterns of Secondary Genes in CGN of PSMG4 in PRAD** reveals significantly switched correlations in cancer and normal tissues in genes involved in the Ferroptosis pathway.

| Core Gene | Secondary Gene | $\mathrm{Corr}_{\mathrm{tumor}}$ | p-value ($\mathrm{Corr}_{\mathrm{tumor}}$) | $\mathrm{Corr}_{\mathrm{normal}}$ | p-value ($\mathrm{Corr}_{\mathrm{normal}}$) | $\lvert Corr_{tumor} - Corr_{normal} \rvert$ |
|---|---|---|---|---|---|---|
| PSMG4 | ACSL4 | -0.165 | 4.0E-04 | 0.618 | 1.7E-06 | 0.783 |
| PSMG4 | MAP1LC3B | 0.040 | 4.0E-01 | 0.767 | 8.4E-11 | 0.727 |
| PSMG4 | ATG5 | 0.033 | 4.7E-01 | 0.753 | 2.9E-10 | 0.719 |
| PSMG4 | PRNP | -0.100 | 3.2E-02 | 0.602 | 3.7E-06 | 0.702 |
| PSMG4 | NCOA4 | -0.007 | 8.8E-01 | 0.687 | 3.6E-08 | 0.694 |
| PSMG4 | PCBP1 | 0.099 | 3.3E-02 | 0.776 | 3.8E-11 | 0.676 |
| PSMG4 | LPCAT3 | 0.015 | 7.5E-01 | 0.688 | 3.5E-08 | 0.673 |
| PSMG4 | VDAC3 | 0.151 | 1.2E-03 | 0.821 | 2.8E-13 | 0.670 |
| PSMG4 | FTH1 | 0.158 | 6.9E-04 | 0.812 | 8.8E-13 | 0.653 |
| PSMG4 | SLC39A14 | 0.007 | 8.9E-01 | 0.652 | 2.9E-07 | 0.645 |
| PSMG4 | SLC40A1 | -0.086 | 6.5E-02 | 0.541 | 5.1E-05 | 0.627 |
| PSMG4 | SLC11A2 | 0.107 | 2.2E-02 | 0.730 | 1.7E-09 | 0.623 |
| PSMG4 | PCBP2 | 0.176 | 1.6E-04 | 0.776 | 3.5E-11 | 0.601 |
| PSMG4 | VDAC2 | 0.239 | 2.3E-07 | 0.824 | 1.9E-13 | 0.585 |
| PSMG4 | GCLC | 0.053 | 2.6E-01 | 0.636 | 7.1E-07 | 0.583 |
| PSMG4 | ACSL6 | -0.033 | 4.8E-01 | -0.575 | 1.3E-05 | 0.542 |

**Supplementary Table S8 – Detailed breakdown of all genes identified to be enriched in pathways of Core Gene Networks (CGN)**. Pearson correlation coefficients across normal, tumor and all samples as well as the differential co-expression between normal and tumor are included. Critical values obtained for differential co-expression as well as results of the limma-voom pipeline for differential expression for both core genes and the secondary genes are included.
(Refer Supplementary Excel File)

**Supplementary Table S9 – Breakdown of sample sizes across the 6 different cancers chosen from the TCGA database.**

| TCGA Name | Cancer | Primary Tumor | Normal Tissue |
|---|---|---|---|
| BRCA | Breast Invasive Carcinoma | 1,041 | 111 |
| KIRC | Kidney Renal Clear Cell Carcinoma | 480 | 70 |
| LUAD | Lung Adenocarcinoma | 483 | 54 |
| COAD | Colon Adenocarcinoma | 387 | 37 |
| PRAD | Prostate Adenocarcinoma | 458 | 50 |
| THCA | Thyroid Carcinoma | 444 | 53 |

### Supplementary Figures

**Supplementary Figure S1 - DNA replication (hsa03030) and some core gene interactions highlighted by SCOPE**. **A)** Shows differentially expressed core genes (light blue) and their interactions with other genes (grey) in the pathway while **B)** shows similar interactions with non-differentially expressed core genes. Correlations are indicated as edges ranging from red (-1) to blue (+1). Boxplots show the correlations indicated in the same pathways to highlight differences in distributions along with the p-value for the Kolmogorov-Smirnov test with the null hypothesis being that the two samples (correlations of Tumour and Normal tissues) were drawn from the same distribution.

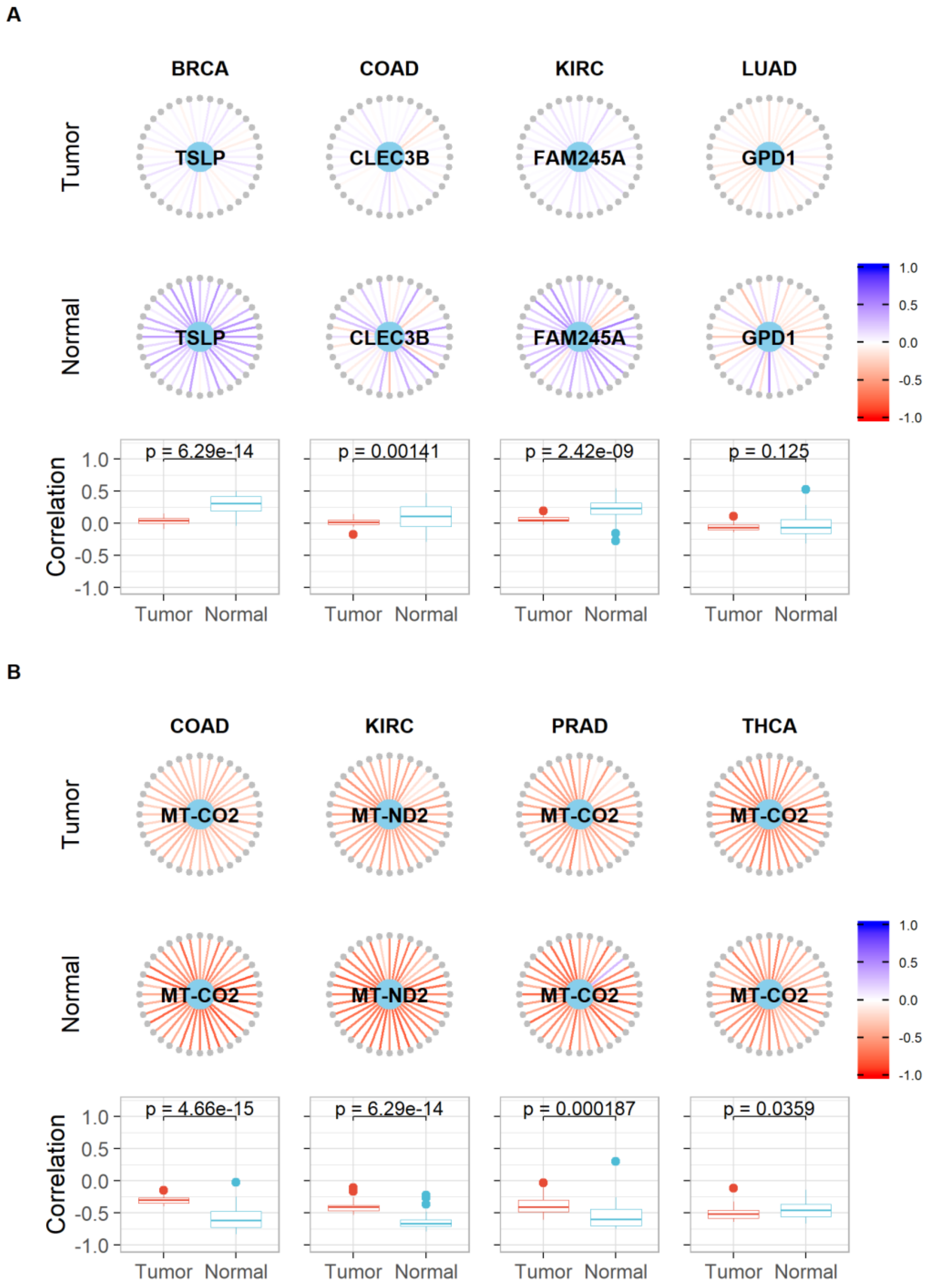

**Supplementary Figure S2 - Proteasome (hsa03050) and some core gene interactions highlighted by SCOPE. A)** Shows differentially expressed core genes (light blue) and their interactions with other genes (grey) in the pathway while **B)** shows similar interactions with non-differentially expressed core genes. Correlations are indicated as edges ranging from red (-1) to blue (+1). Boxplots show the correlations indicated in the same pathways to highlight differences in distributions along with the p-value for the Kolmogorov-Smirnov test with the null hypothesis being that the two samples (correlations of Tumour and Normal tissues) were drawn from the same distribution.

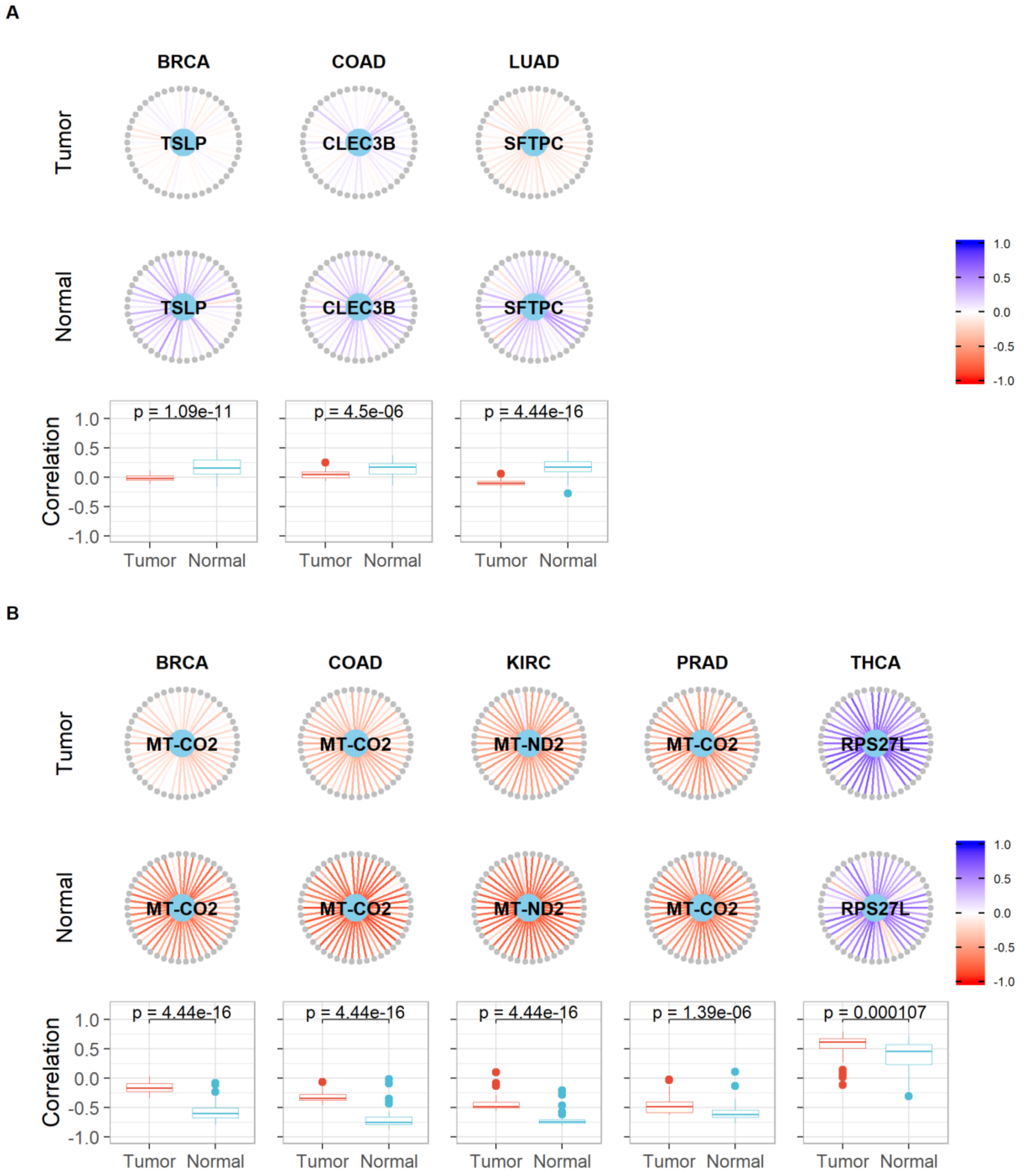

**Supplementary Figure S3 - Spliceosome (hsa03040) and some core gene interactions highlighted by SCOPE. A)** Shows differentially expressed core genes (light blue) and their interactions with other genes (grey) in the pathway while **B)** shows similar interactions with non-differentially expressed core genes. Correlations are indicated as edges ranging from red (-1) to blue (+1). Boxplots show the correlations indicated in the same pathways to highlight differences in distributions along with the p-value for the Kolmogorov-Smirnov test with the null hypothesis being that the two samples (correlations of Tumour and Normal tissues) were drawn from the same distribution.

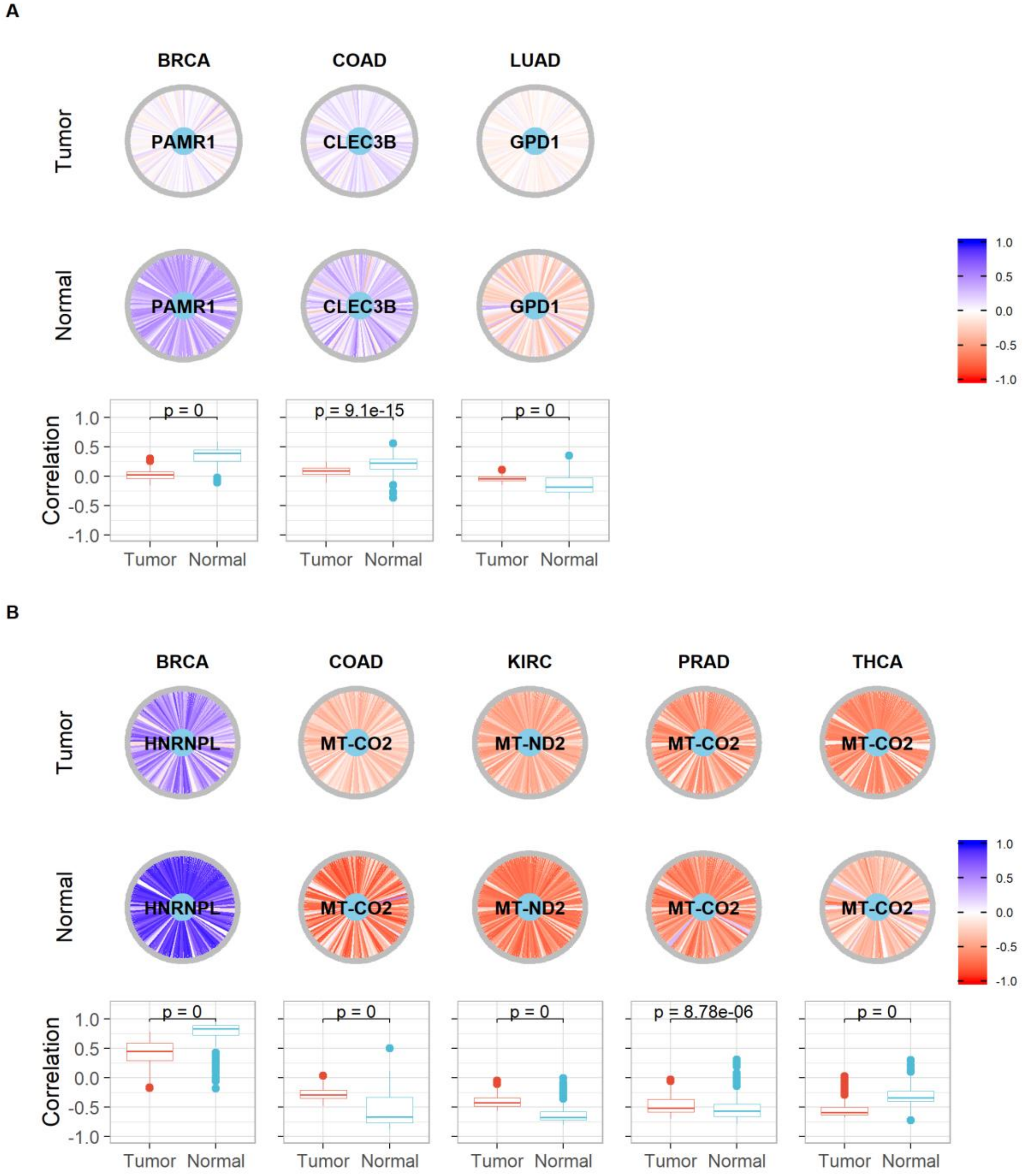

**Supplementary Figure S4 - RNA polymerase (hsa03020) and some core gene interactions highlighted by SCOPE. A)** Shows differentially expressed core genes (light blue) and their interactions with other genes (grey) in the pathway while **B)** shows similar interactions with non-differentially expressed core genes. Correlations are indicated as edges ranging from red (-1) to blue (+1). Boxplots show the correlations indicated in the same pathways to highlight differences in distributions along with the p-value for the Kolmogorov-Smirnov test with the null hypothesis being that the two samples (correlations of Tumour and Normal tissues) were drawn from the same distribution.

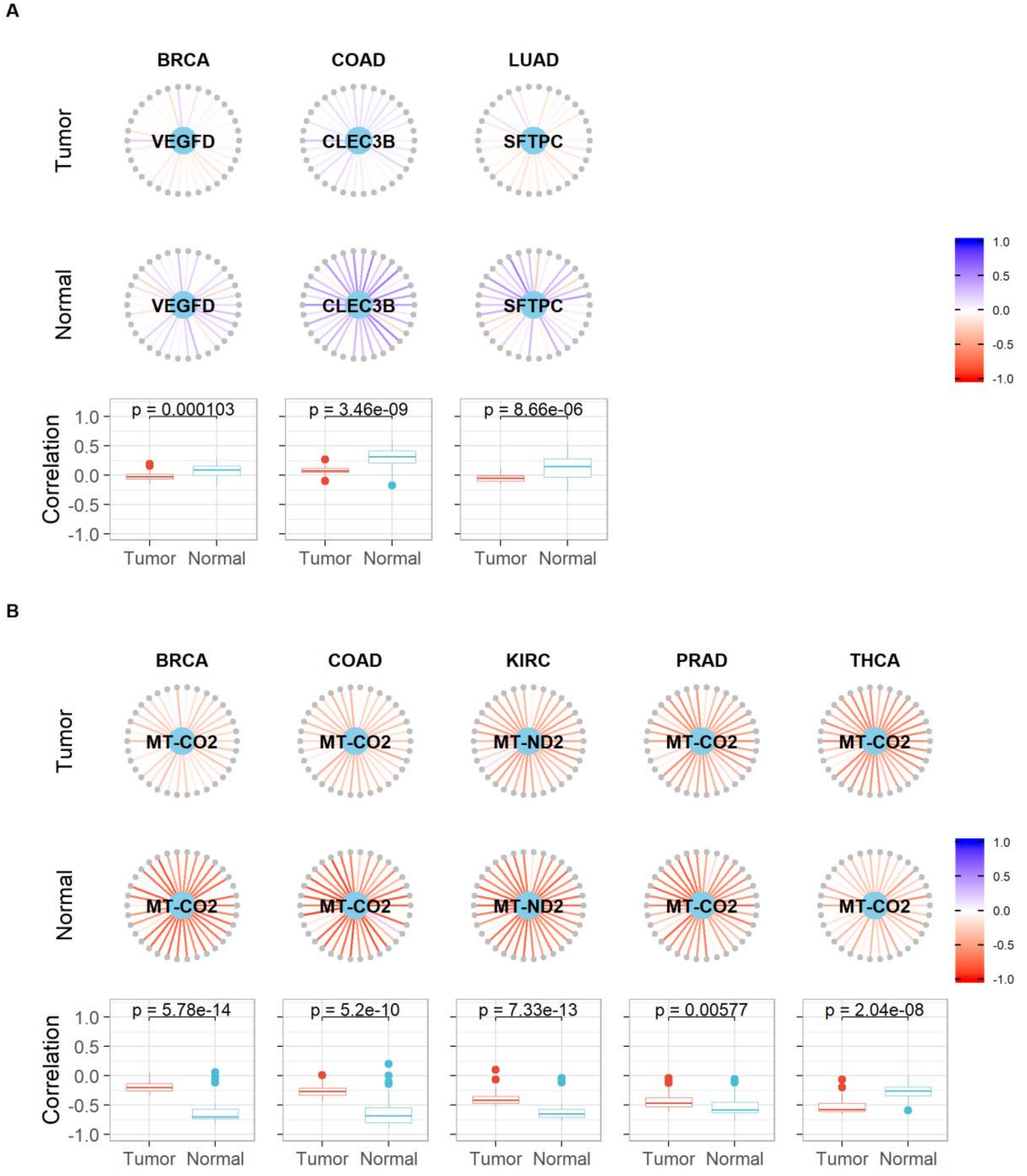

**Supplementary Figure S5 - Base excision repair (hsa03410) and some core gene interactions highlighted by SCOPE. A)** Shows differentially expressed core genes (light blue) and their interactions with other genes (grey) in the pathway while **B)** shows similar interactions with non-differentially expressed core genes. Correlations are indicated as edges ranging from red (-1) to blue (+1). Boxplots show the correlations indicated in the same pathways to highlight differences in distributions along with the p-value for the Kolmogorov-Smirnov test with the null hypothesis being that the two samples (correlations of Tumour and Normal tissues) were drawn from the same distribution.

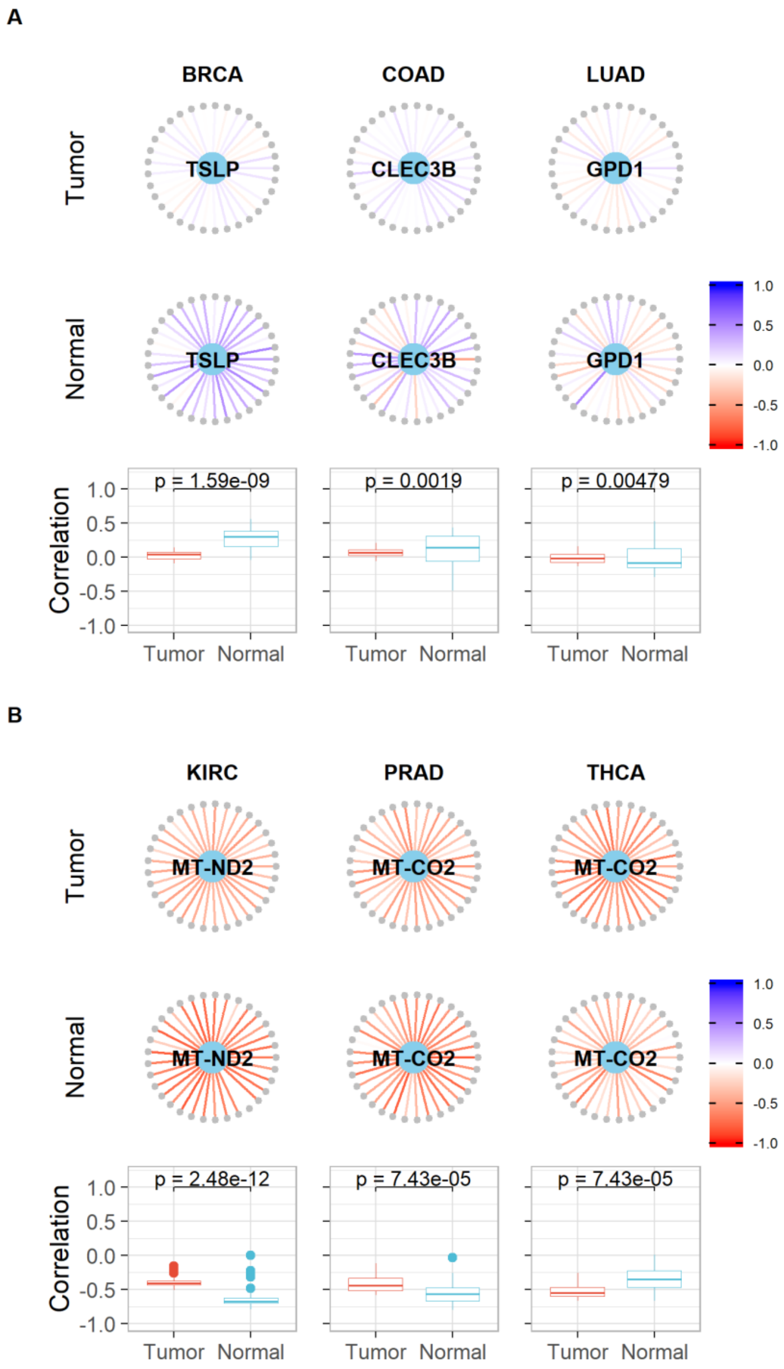

**Supplementary Figure S6 - N-Glycan biosynthesis (hsa00510) and some core gene interactions highlighted by SCOPE. A)** Shows differentially expressed core genes (light blue) and their interactions with other genes (grey) in the pathway while **B)** shows similar interactions with non-differentially expressed core genes. Correlations are indicated as edges ranging from red (-1) to blue (+1). Boxplots show the correlations indicated in the same pathways to highlight differences in distributions along with the p-value for the Kolmogorov-Smirnov test with the null hypothesis being that the two samples (correlations of Tumour and Normal tissues) were drawn from the same distribution.

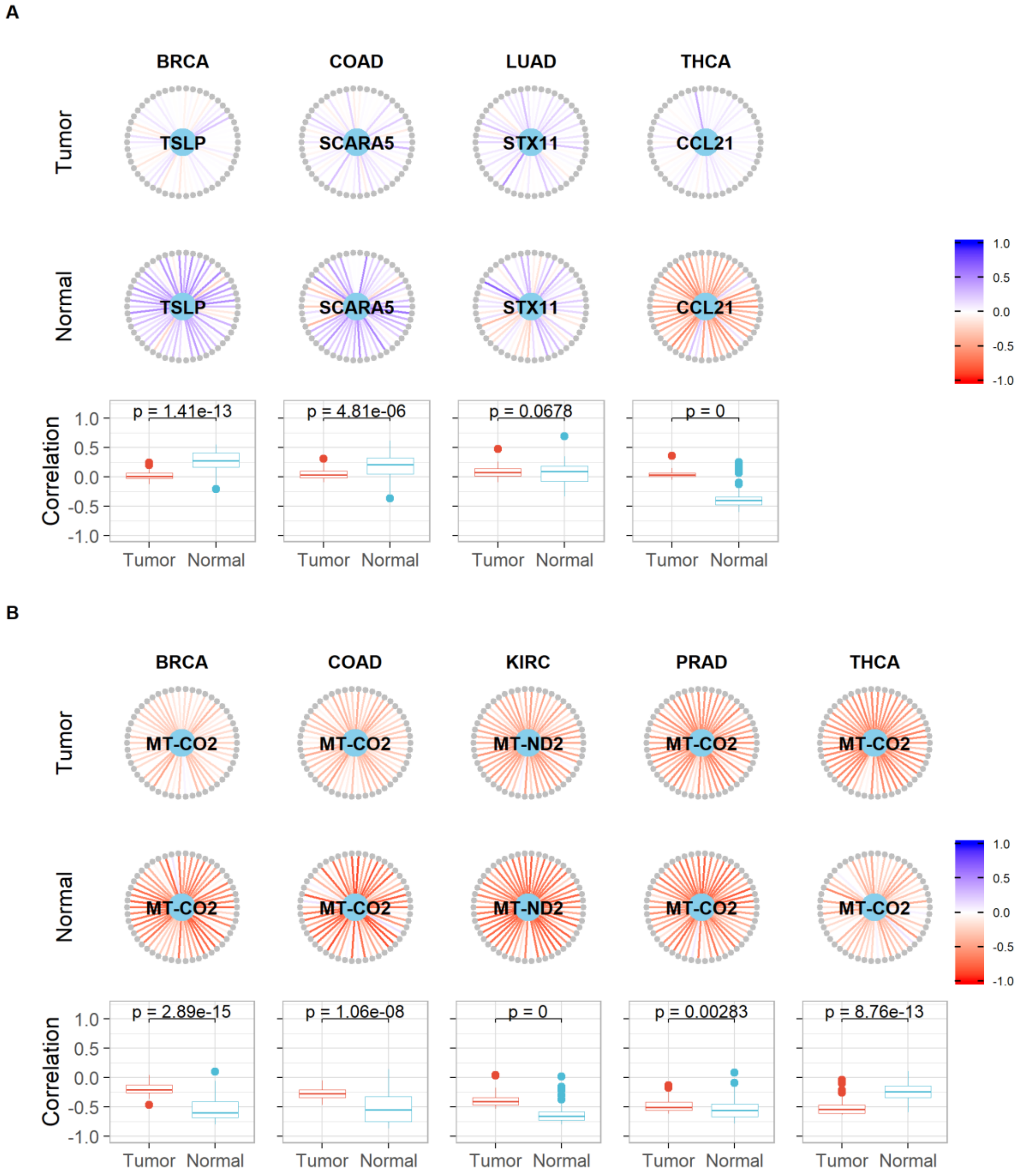

**Supplementary Figure S7 - Fanconi anemia pathway (hsa03460) and some core gene interactions highlighted by SCOPE. A)** Shows differentially expressed core genes (light blue) and their interactions with other genes (grey) in the pathway while **B)** shows similar interactions with non-differentially expressed core genes. Correlations are indicated as edges ranging from red (-1) to blue (+1). Boxplots show the correlations indicated in the same pathways to highlight differences in distributions along with the p-value for the Kolmogorov-Smirnov test with the null hypothesis being that the two samples (correlations of Tumour and Normal tissues) were drawn from the same distribution.

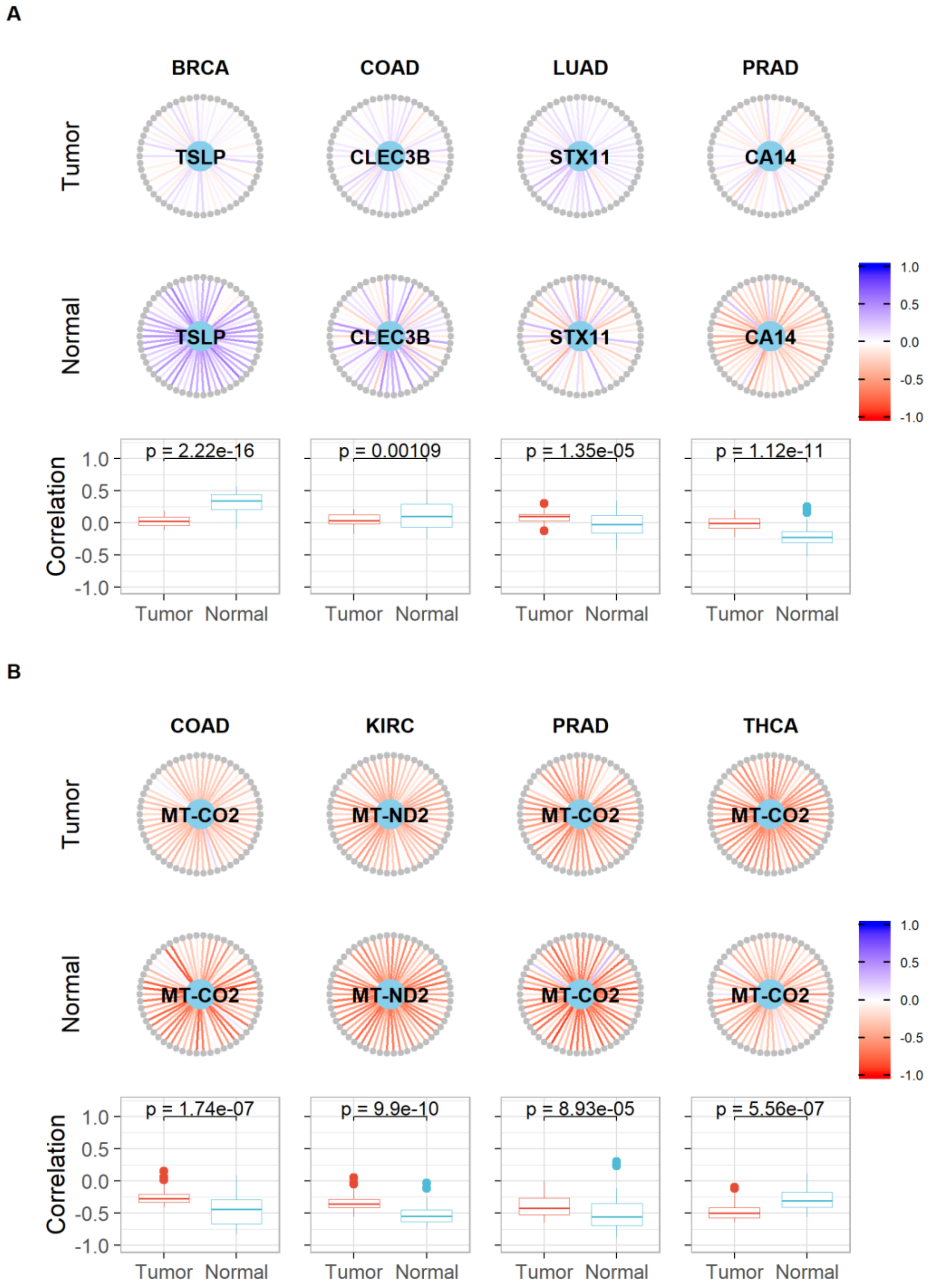

**Supplementary Figure S8 - Cell cycle (hsa04110) and some core gene interactions highlighted by SCOPE. A)** Shows differentially expressed core genes (light blue) and their interactions with other genes (grey) in the pathway while **B)** shows similar interactions with non-differentially expressed core genes. Correlations are indicated as edges ranging from red (-1) to blue (+1). Boxplots show the correlations indicated in the same pathways to highlight differences in distributions along with the p-value for the Kolmogorov-Smirnov test with the null hypothesis being that the two samples (correlations of Tumour and Normal tissues) were drawn from the same distribution.

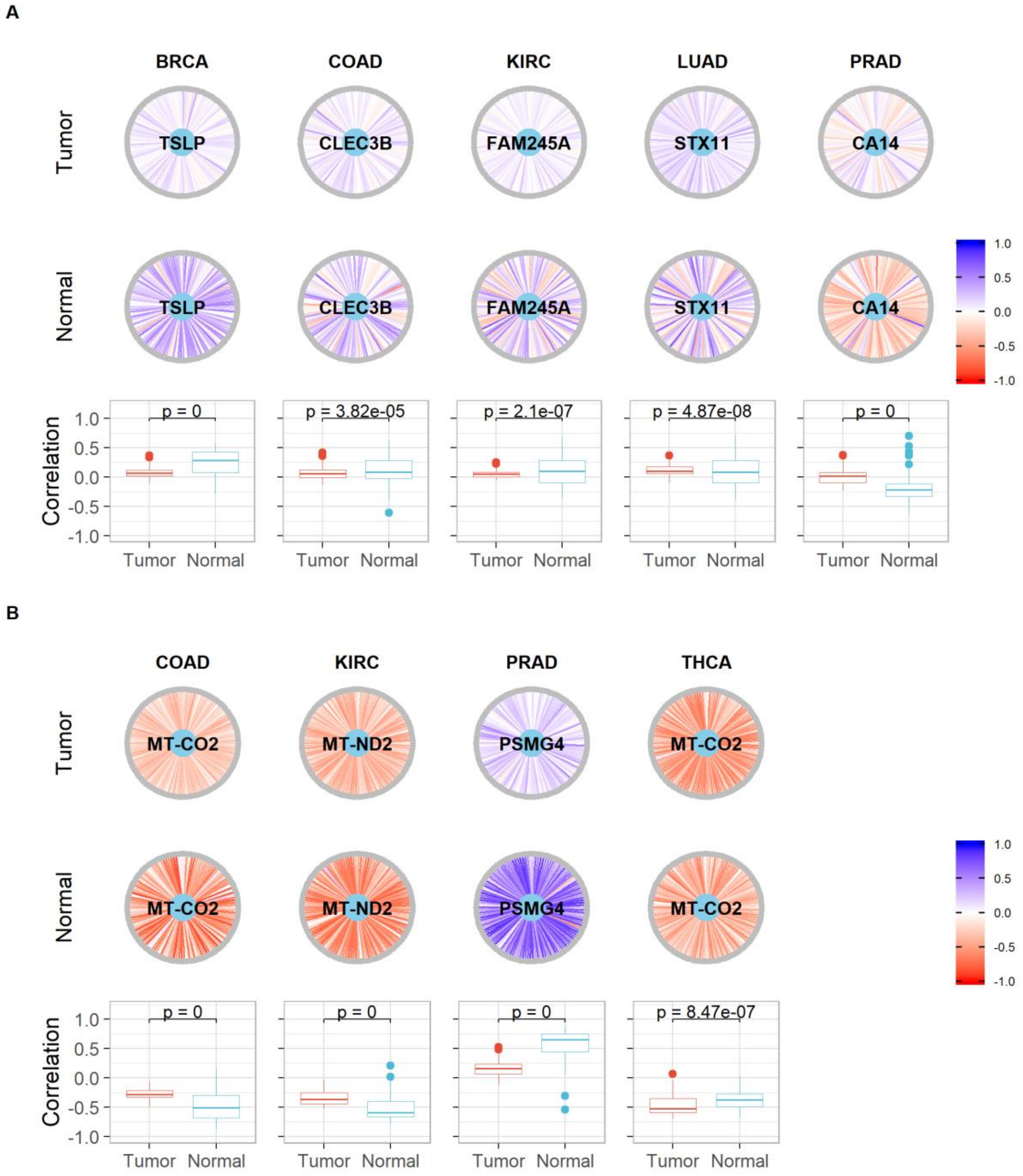

**Supplementary Figure S9 - Protein processing in endoplasmic reticulum (hsa04141) and some core gene interactions highlighted by SCOPE**. **A)** Shows differentially expressed core genes (light blue) and their interactions with other genes (grey) in the pathway while **B)** shows similar interactions with non-differentially expressed core genes. Correlations are indicated as edges ranging from red (-1) to blue (+1). Boxplots show the correlations indicated in the same pathways to highlight differences in distributions along with the p-value for the Kolmogorov-Smirnov test with the null hypothesis being that the two samples (correlations of Tumour and Normal tissues) were drawn from the same distribution.

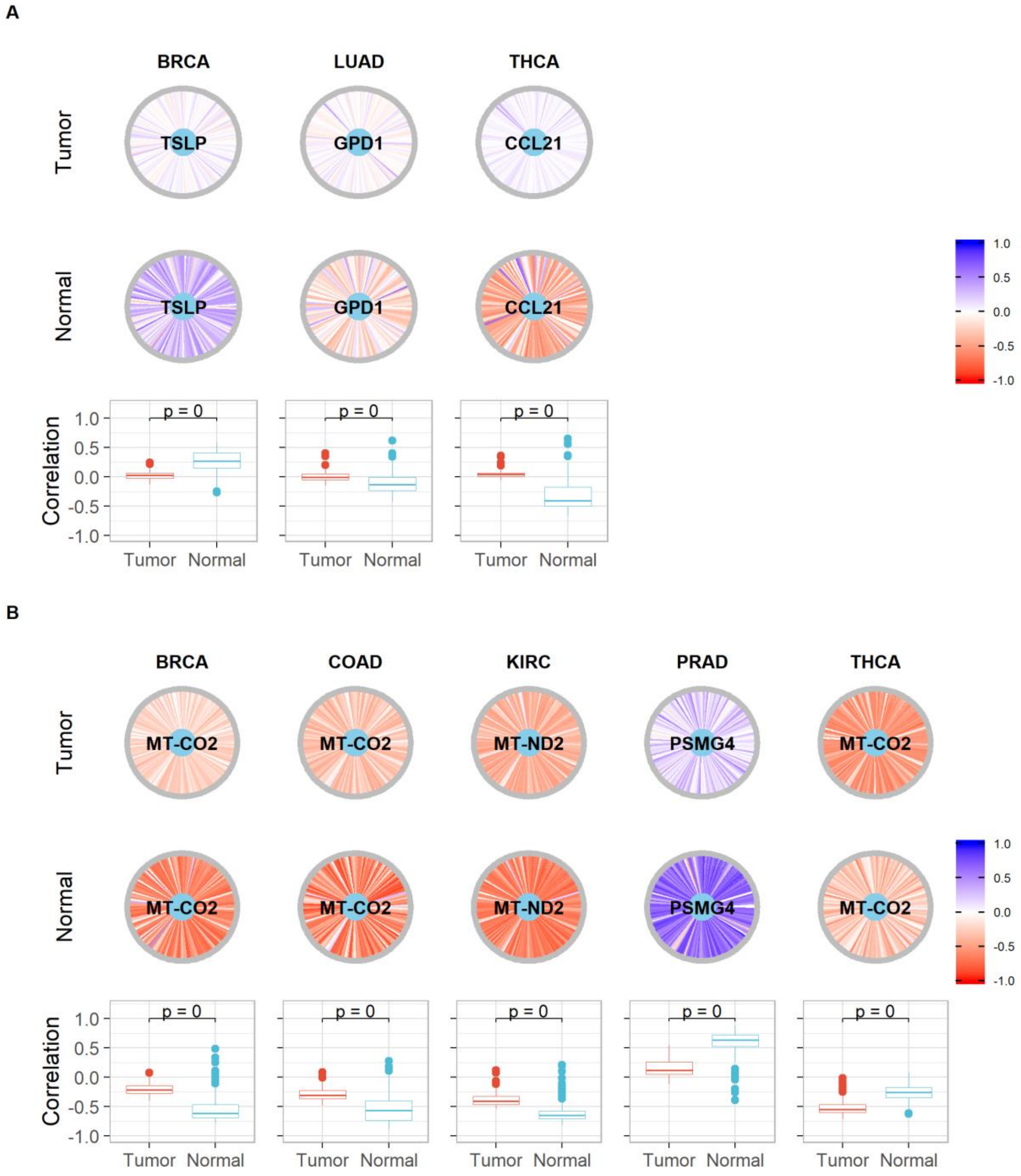

**Supplementary Figure S10 - RNA degradation (hsa03018) and some core gene interactions highlighted by SCOPE. A)** Shows differentially expressed core genes (light blue) and their interactions with other genes (grey) in the pathway while **B)** shows similar interactions with non-differentially expressed core genes. Correlations are indicated as edges ranging from red (-1) to blue (+1). Boxplots show the correlations indicated in the same pathways to highlight differences in distributions along with the p-value for the Kolmogorov-Smirnov test with the null hypothesis being that the two samples (correlations of Tumour and Normal tissues) were drawn from the same distribution.

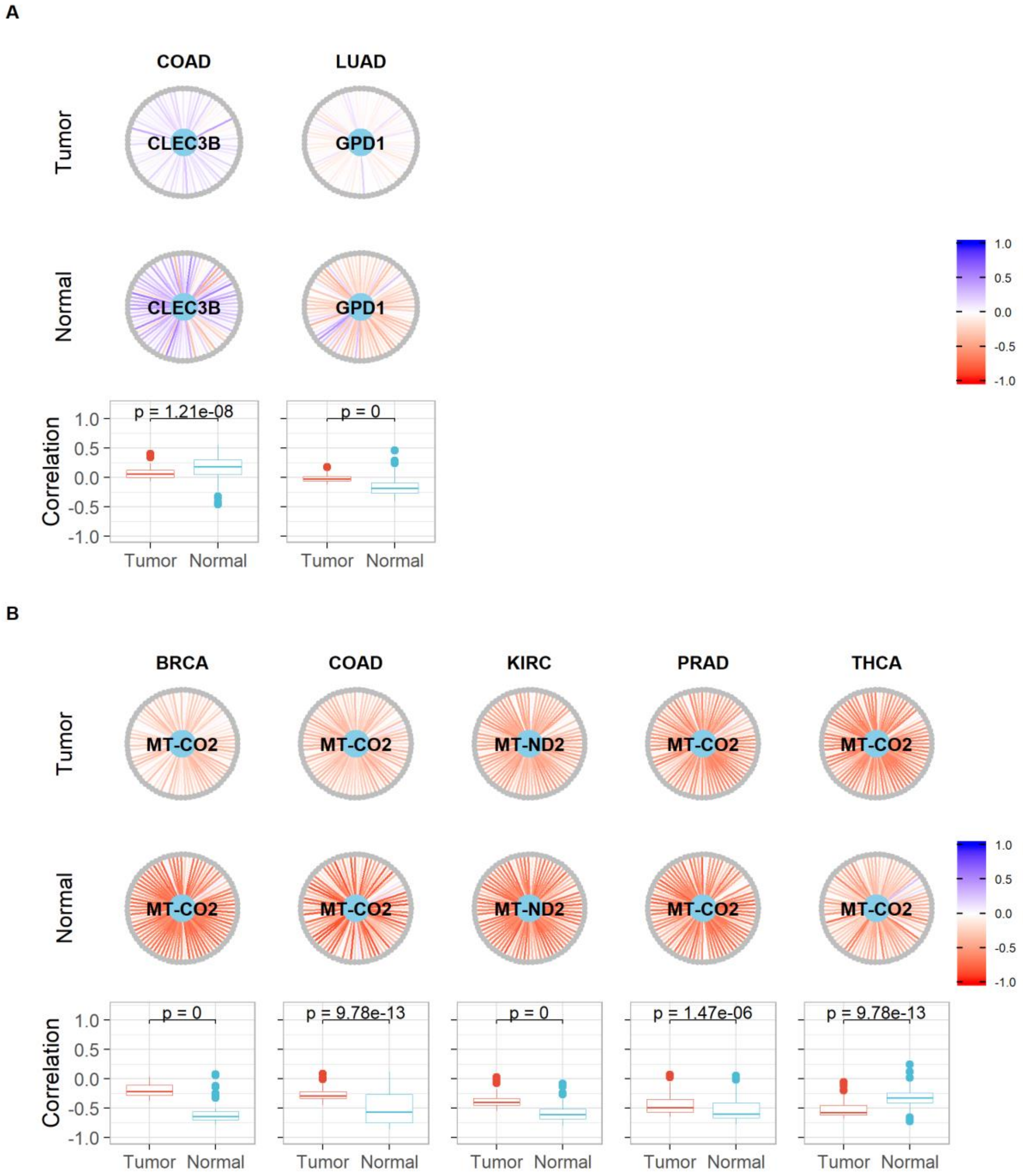

**Supplementary Figure S11 - Homologous recombination (hsa03440) and some core gene interactions highlighted by SCOPE. A)** Shows differentially expressed core genes (light blue) and their interactions with other genes (grey) in the pathway while **B)** shows similar interactions with non-differentially expressed core genes. Correlations are indicated as edges ranging from red (-1) to blue (+1). Boxplots show the correlations indicated in the same pathways to highlight differences in distributions along with the p-value for the Kolmogorov-Smirnov test with the null hypothesis being that the two samples (correlations of Tumour and Normal tissues) were drawn from the same distribution.

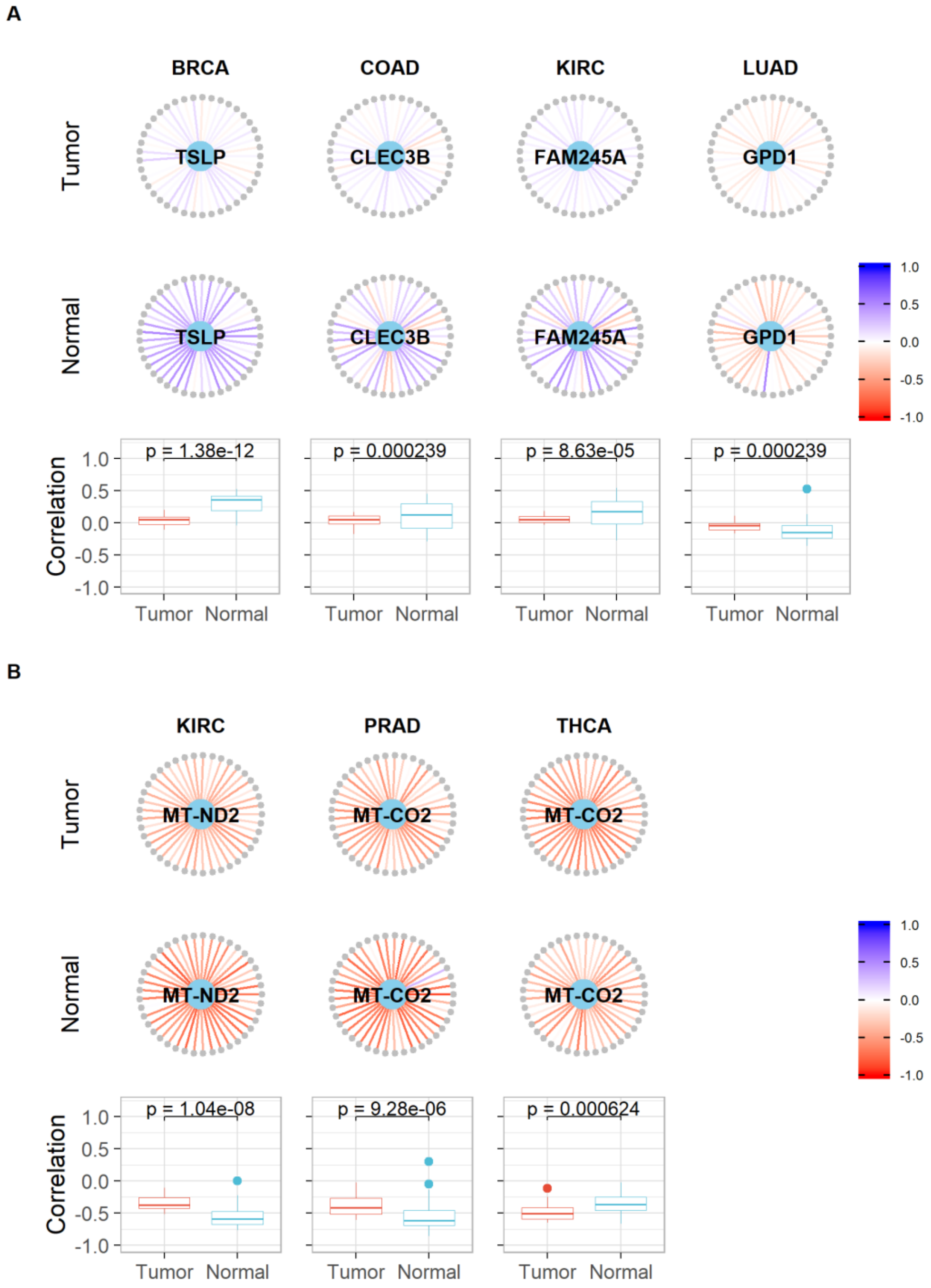

**Supplementary Figure S12 - RNA transport (hsa03013) and some core gene interactions highlighted by SCOPE. A)** Shows differentially expressed core genes (light blue) and their interactions with other genes (grey) in the pathway while **B)** shows similar interactions with non-differentially expressed core genes. Correlations are indicated as edges ranging from red (-1) to blue (+1). Boxplots show the correlations indicated in the same pathways to highlight differences in distributions along with the p-value for the Kolmogorov-Smirnov test with the null hypothesis being that the two samples (correlations of Tumour and Normal tissues) were drawn from the same distribution.

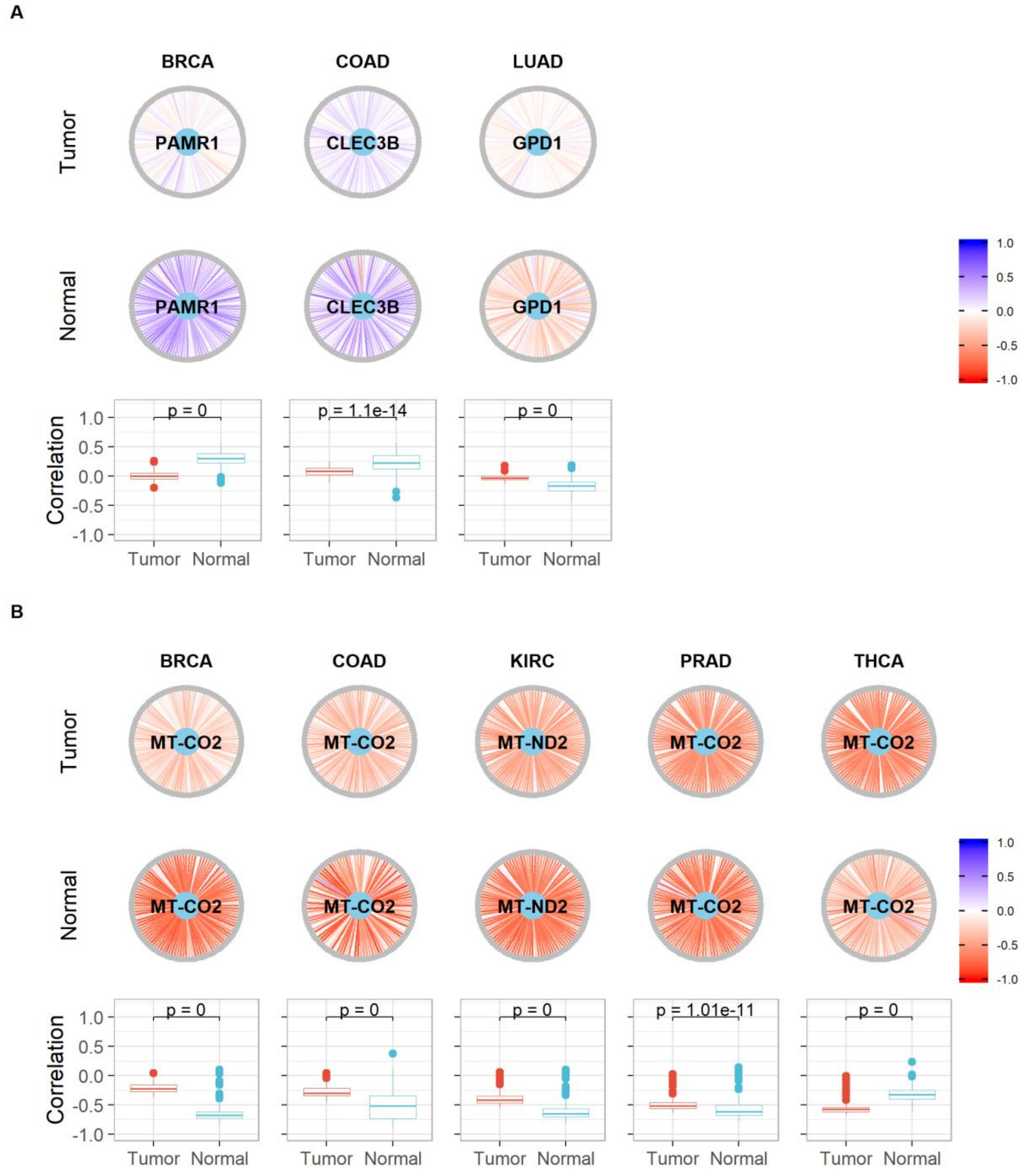

**Supplementary Figure S13 - mRNA surveillance pathway (hsa03015) and some core gene interactions highlighted by SCOPE. A)** Shows differentially expressed core genes (light blue) and their interactions with other genes (grey) in the pathway while **B)** shows similar interactions with non-differentially expressed core genes. Correlations are indicated as edges ranging from red (-1) to blue (+1). Boxplots show the correlations indicated in the same pathways to highlight differences in distributions along with the p-value for the Kolmogorov-Smirnov test with the null hypothesis being that the two samples (correlations of Tumour and Normal tissues) were drawn from the same distribution.

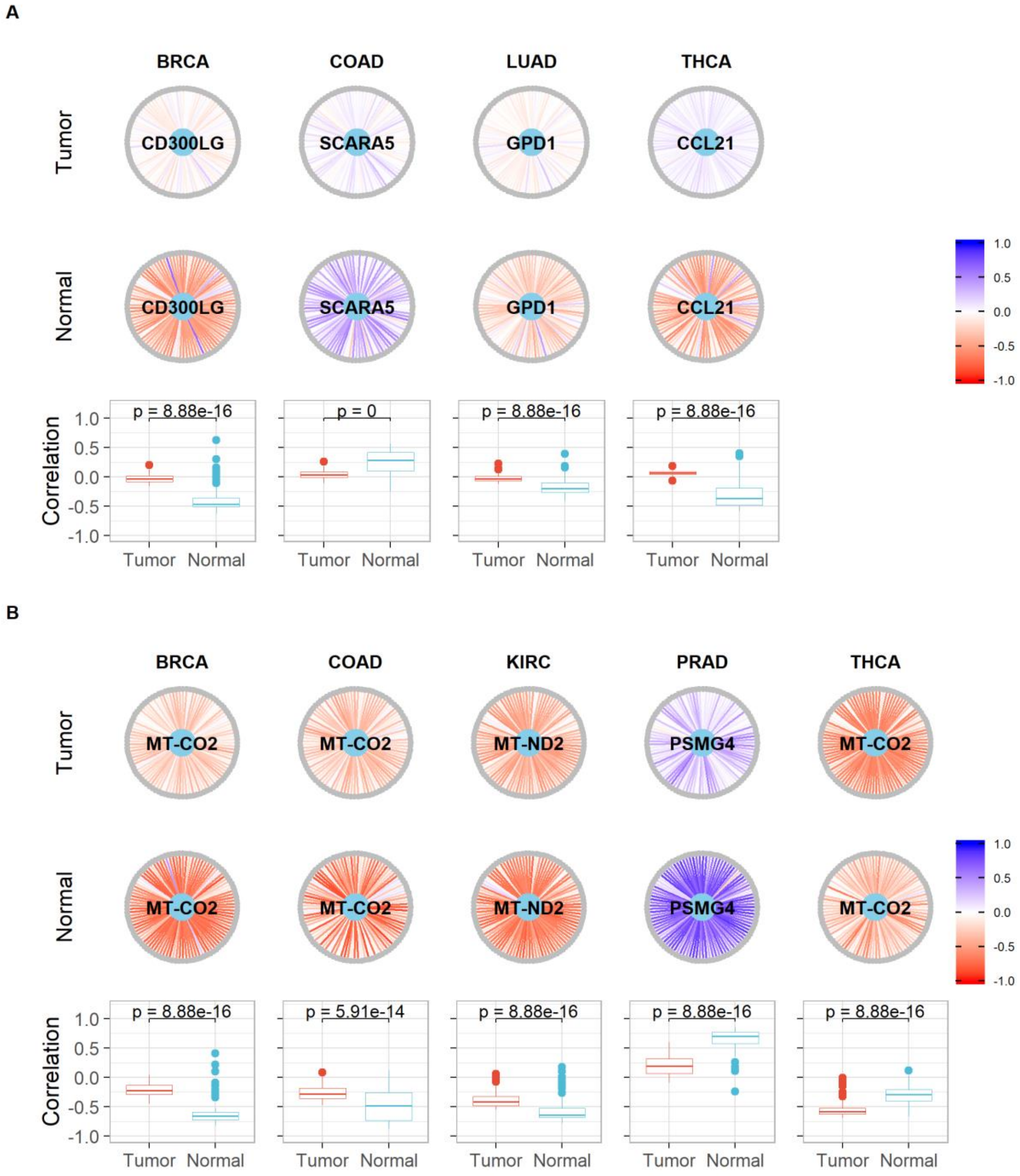

**Supplementary Figure S14 - Pyrimidine metabolism (hsa00240) and some core gene interactions highlighted by SCOPE**. **A)** Shows differentially expressed core genes (light blue) and their interactions with other genes (grey) in the pathway while **B)** shows similar interactions with non-differentially expressed core genes. Correlations are indicated as edges ranging from red (-1) to blue (+1). Boxplots show the correlations indicated in the same pathways to highlight differences in distributions along with the p-value for the Kolmogorov-Smirnov test with the null hypothesis being that the two samples (correlations of Tumour and Normal tissues) were drawn from the same distribution.

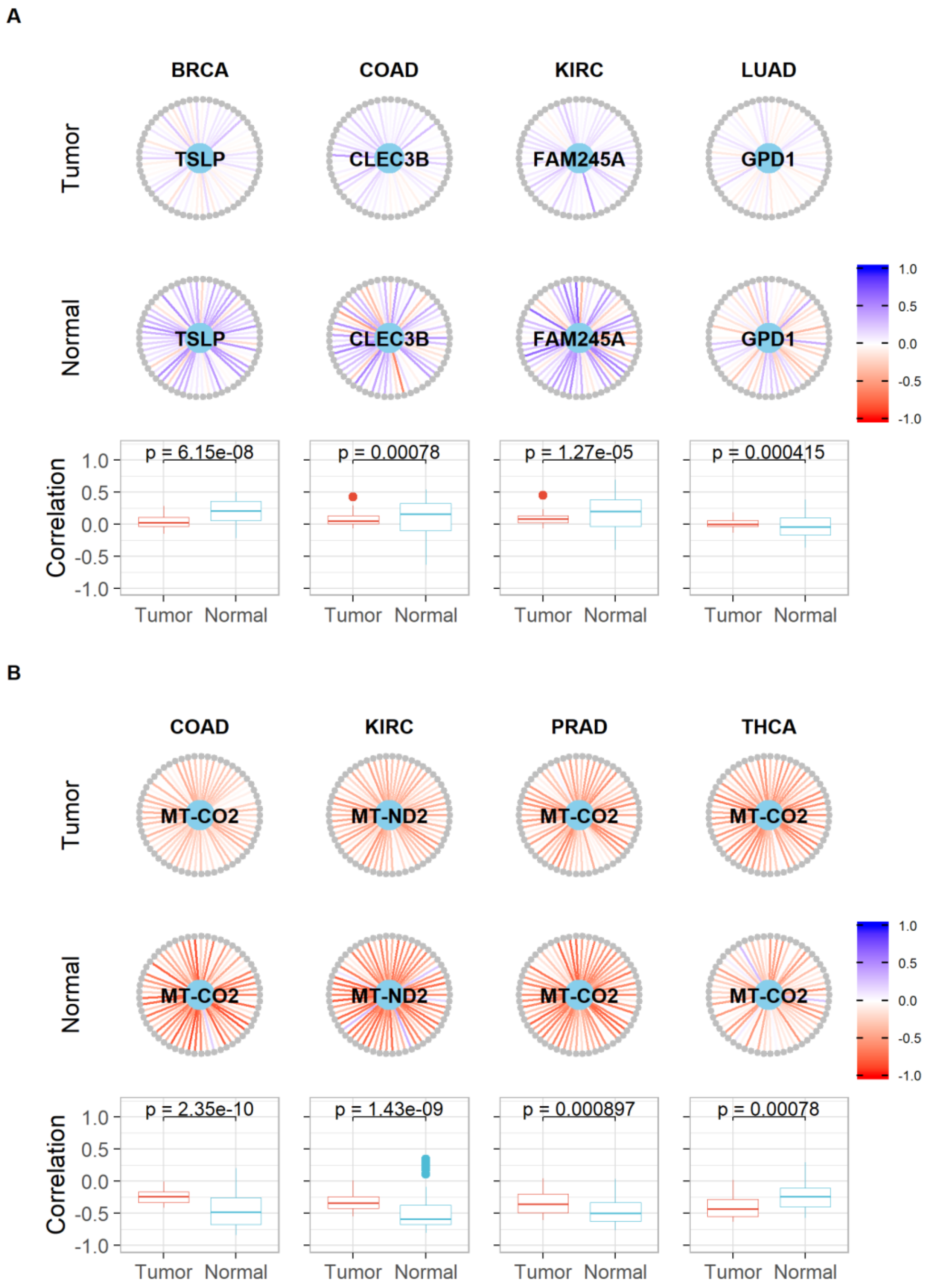

**Supplementary Figure S15 - Thermogenesis (hsa04714) and some core gene interactions highlighted by SCOPE. A)** Shows differentially expressed core genes (light blue) and their interactions with other genes (grey) in the pathway while **B)** shows similar interactions with non-differentially expressed core genes. Correlations are indicated as edges ranging from red (-1) to blue (+1). Boxplots show the correlations indicated in the same pathways to highlight differences in distributions along with the p-value for the Kolmogorov-Smirnov test with the null hypothesis being that the two samples (correlations of Tumour and Normal tissues) were drawn from the same distribution.

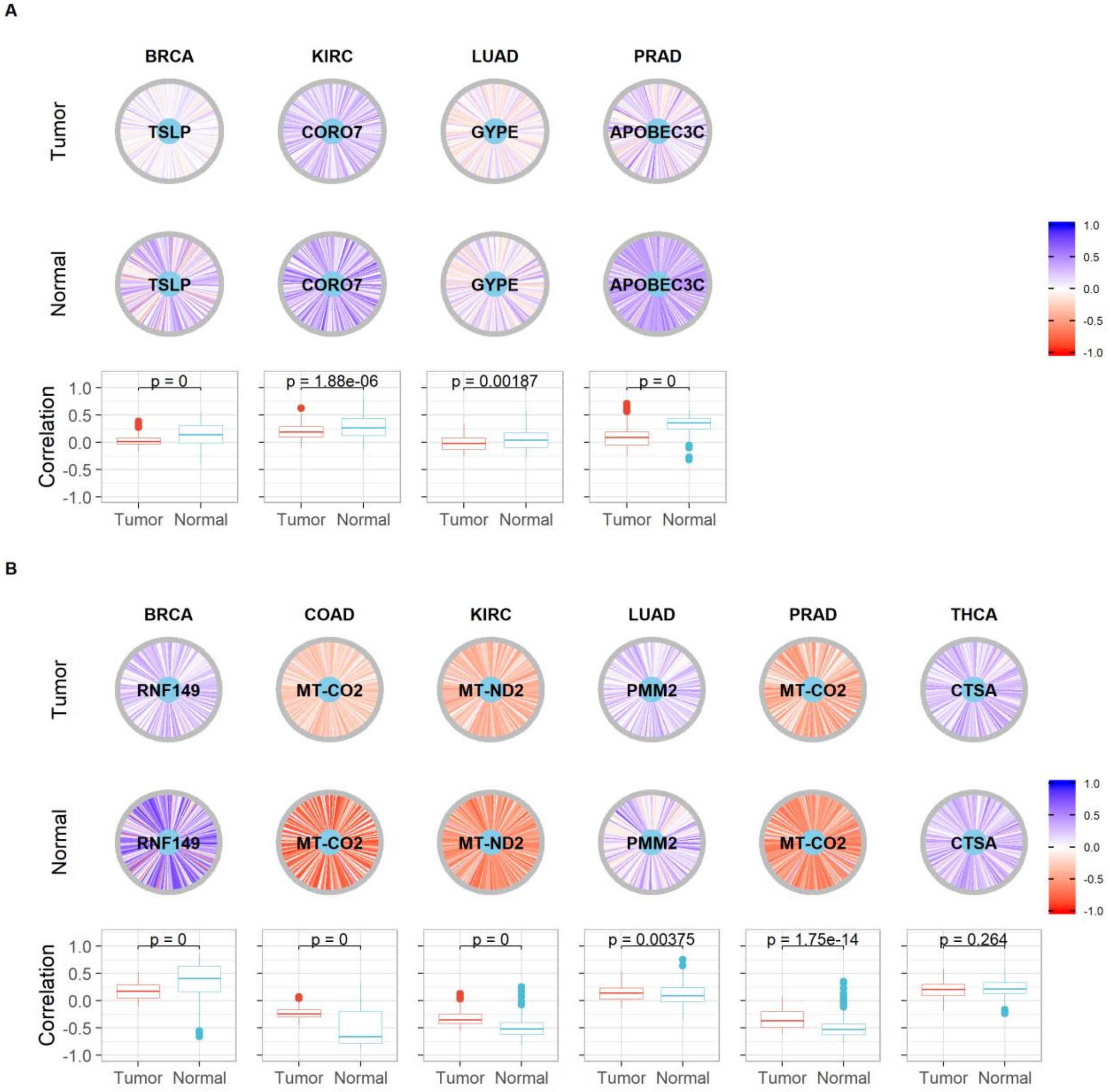

**Supplementary Figure S16 - p53 signaling pathway (hsa04115) and some core gene interactions highlighted by SCOPE. A)** Shows differentially expressed core genes (light blue) and their interactions with other genes (grey) in the pathway while **B)** shows similar interactions with non-differentially expressed core genes. Correlations are indicated as edges ranging from red (-1) to blue (+1). Boxplots show the correlations indicated in the same pathways to highlight differences in distributions along with the p-value for the Kolmogorov-Smirnov test with the null hypothesis being that the two samples (correlations of Tumour and Normal tissues) were drawn from the same distribution.

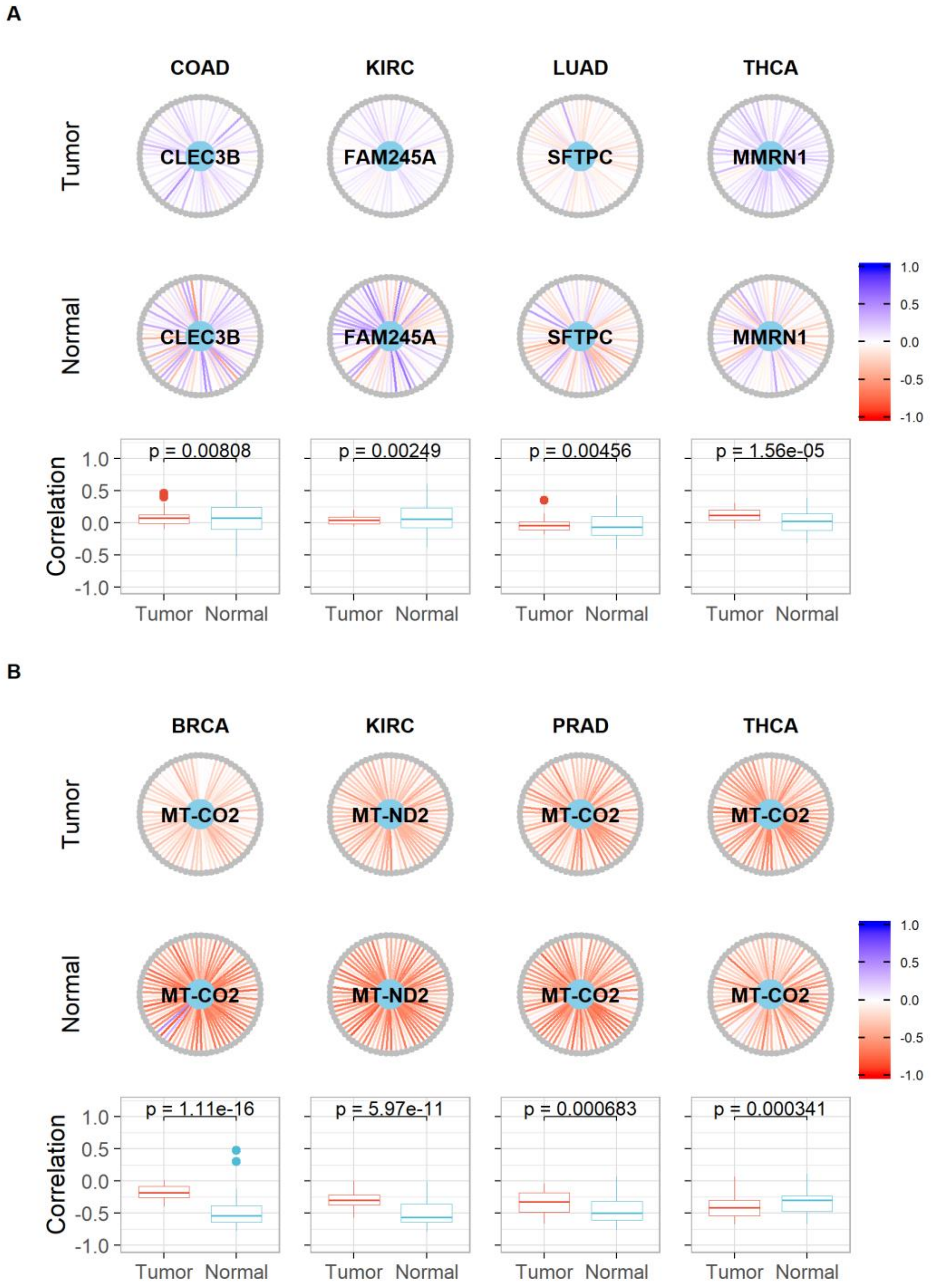

**Supplementary Figure S17 - Alzheimer disease (hsa05010) and some core gene interactions highlighted by SCOPE. A)** Shows differentially expressed core genes (light blue) and their interactions with other genes (grey) in the pathway while **B)** shows similar interactions with non-differentially expressed core genes. Correlations are indicated as edges ranging from red (-1) to blue (+1). Boxplots show the correlations indicated in the same pathways to highlight differences in distributions along with the p-value for the Kolmogorov-Smirnov test with the null hypothesis being that the two samples (correlations of Tumour and Normal tissues) were drawn from the same distribution.

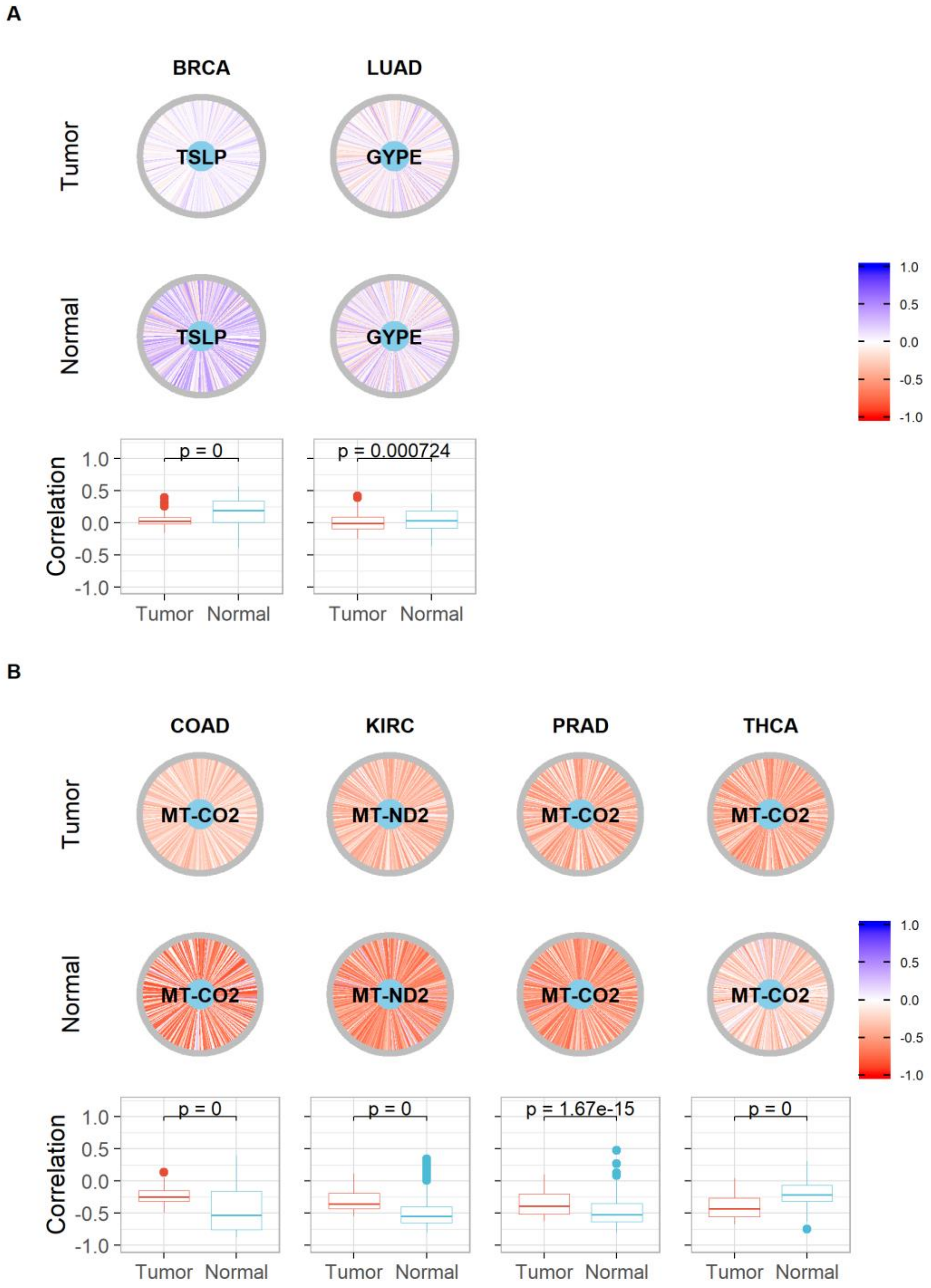

**Supplementary Figure S18 - Ribosome (hsa03010) and some core gene interactions highlighted by SCOPE. A)** Shows differentially expressed core genes (light blue) and their interactions with other genes (grey) in the pathway while **B)** shows similar interactions with non-differentially expressed core genes. Correlations are indicated as edges ranging from red (-1) to blue (+1). Boxplots show the correlations indicated in the same pathways to highlight differences in distributions along with the p-value for the Kolmogorov-Smirnov test with the null hypothesis being that the two samples (correlations of Tumour and Normal tissues) were drawn from the same distribution.

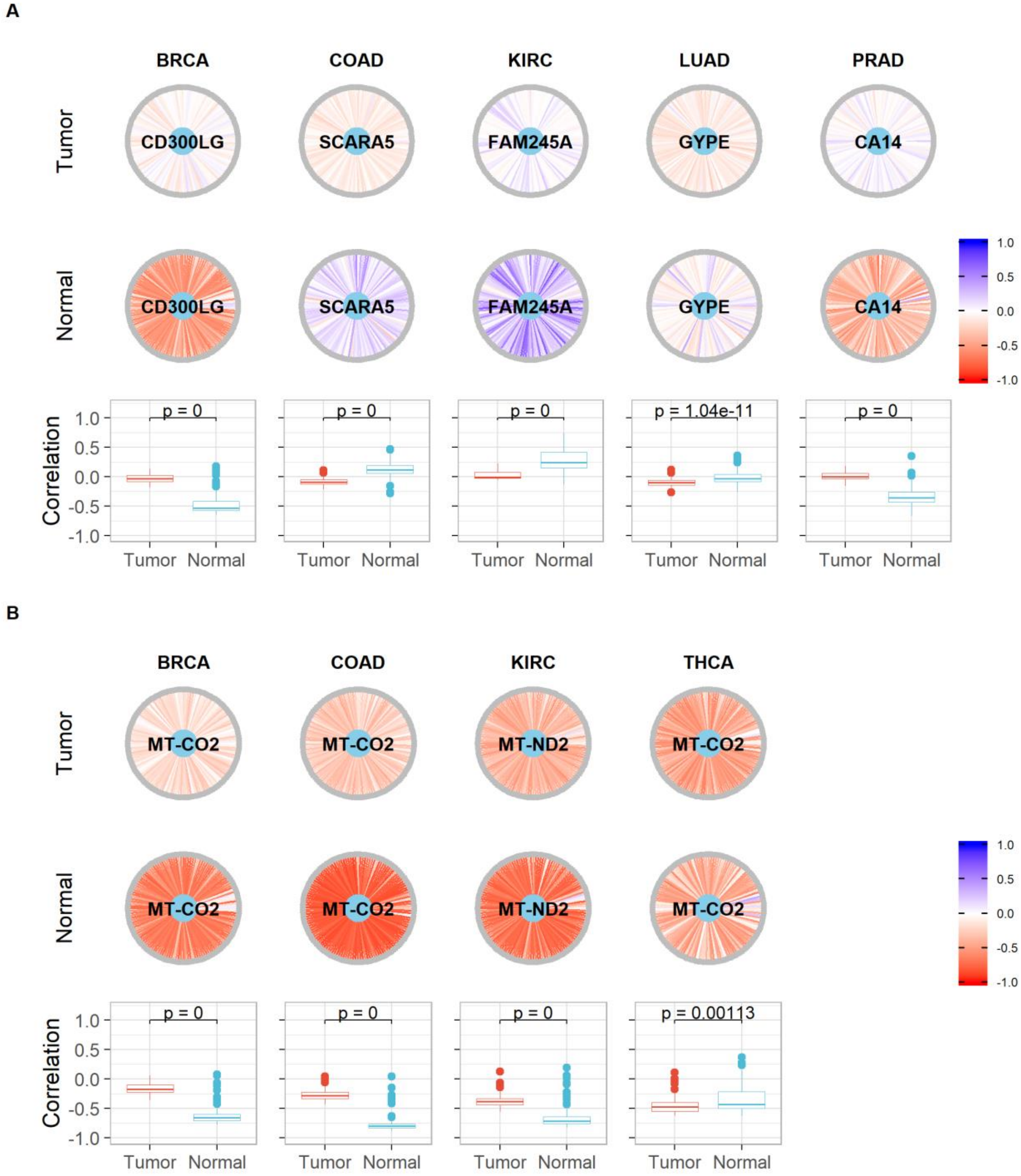

**Supplementary Figure S19 - Oxidative phosphorylation (hsa00190) and some core gene interactions highlighted by SCOPE. A)** Shows differentially expressed core genes (light blue) and their interactions with other genes (grey) in the pathway while **B)** shows similar interactions with non-differentially expressed core genes. Correlations are indicated as edges ranging from red (-1) to blue (+1). Boxplots show the correlations indicated in the same pathways to highlight differences in distributions along with the p-value for the Kolmogorov-Smirnov test with the null hypothesis being that the two samples (correlations of Tumour and Normal tissues) were drawn from the same distribution.

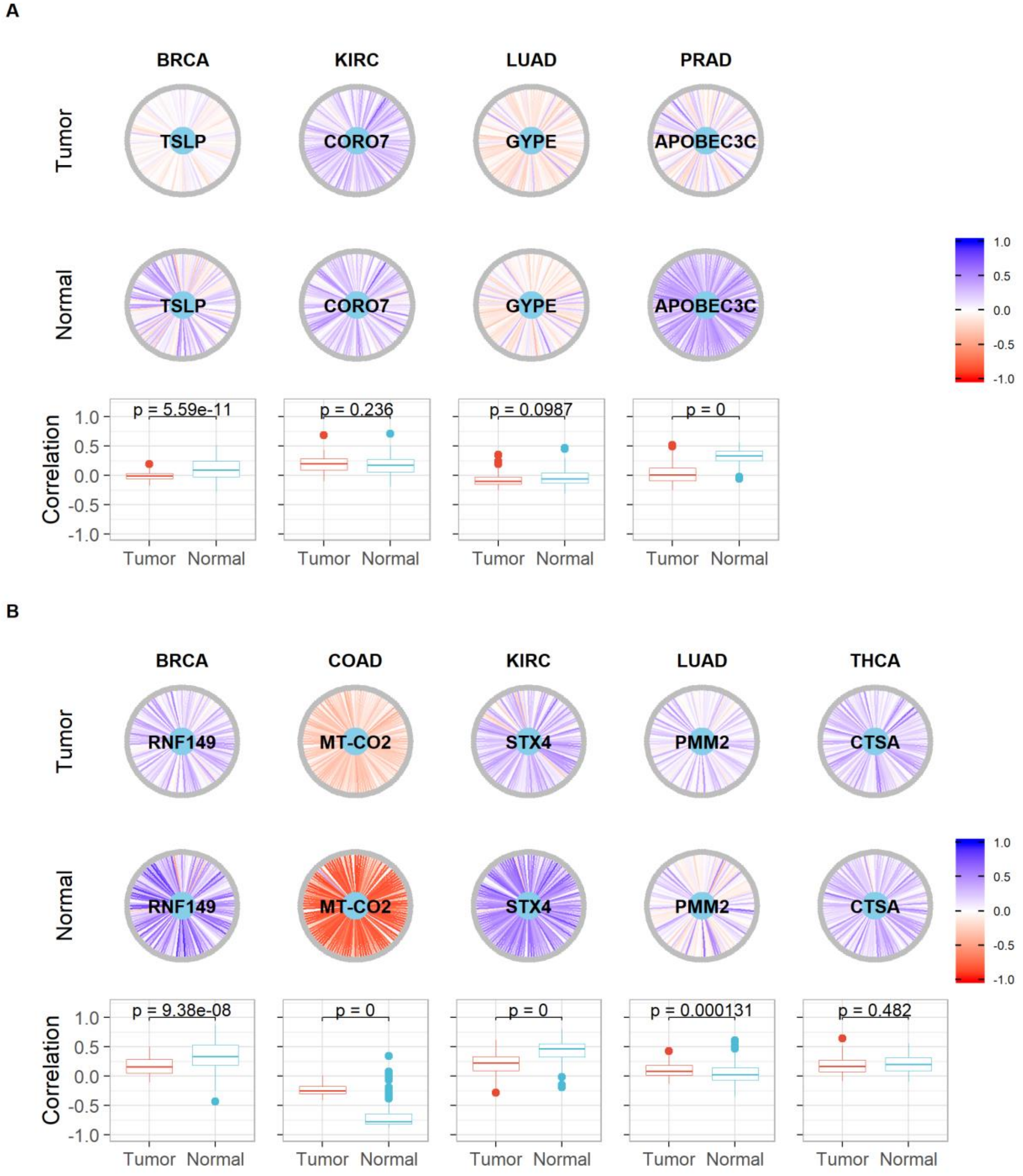

**Supplementary Figure S20 - Parkinson disease (hsa05012) and some core gene interactions highlighted by SCOPE. A)** Shows differentially expressed core genes (light blue) and their interactions with other genes (grey) in the pathway while **B)** shows similar interactions with non-differentially expressed core genes. Correlations are indicated as edges ranging from red (-1) to blue (+1). Boxplots show the correlations indicated in the same pathways to highlight differences in distributions along with the p-value for the Kolmogorov-Smirnov test with the null hypothesis being that the two samples (correlations of Tumour and Normal tissues) were drawn from the same distribution.

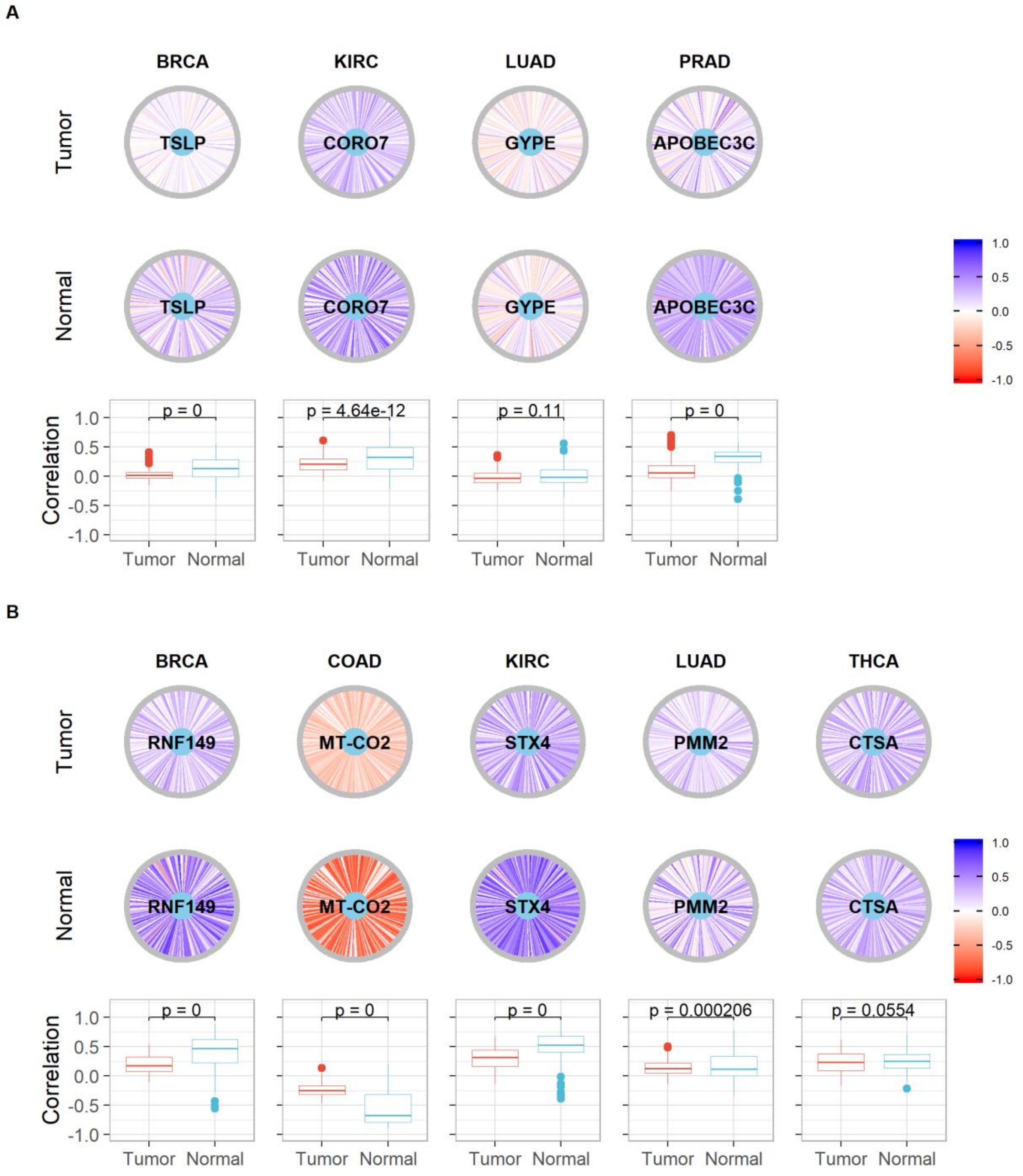

**Supplementary Figure S21 - Purine metabolism (hsa00230) and some core gene interactions highlighted by SCOPE. A)** Shows differentially expressed core genes (light blue) and their interactions with other genes (grey) in the pathway while **B)** shows similar interactions with non-differentially expressed core genes. Correlations are indicated as edges ranging from red (-1) to blue (+1). Boxplots show the correlations indicated in the same pathways to highlight differences in distributions along with the p-value for the Kolmogorov-Smirnov test with the null hypothesis being that the two samples (correlations of Tumour and Normal tissues) were drawn from the same distribution.

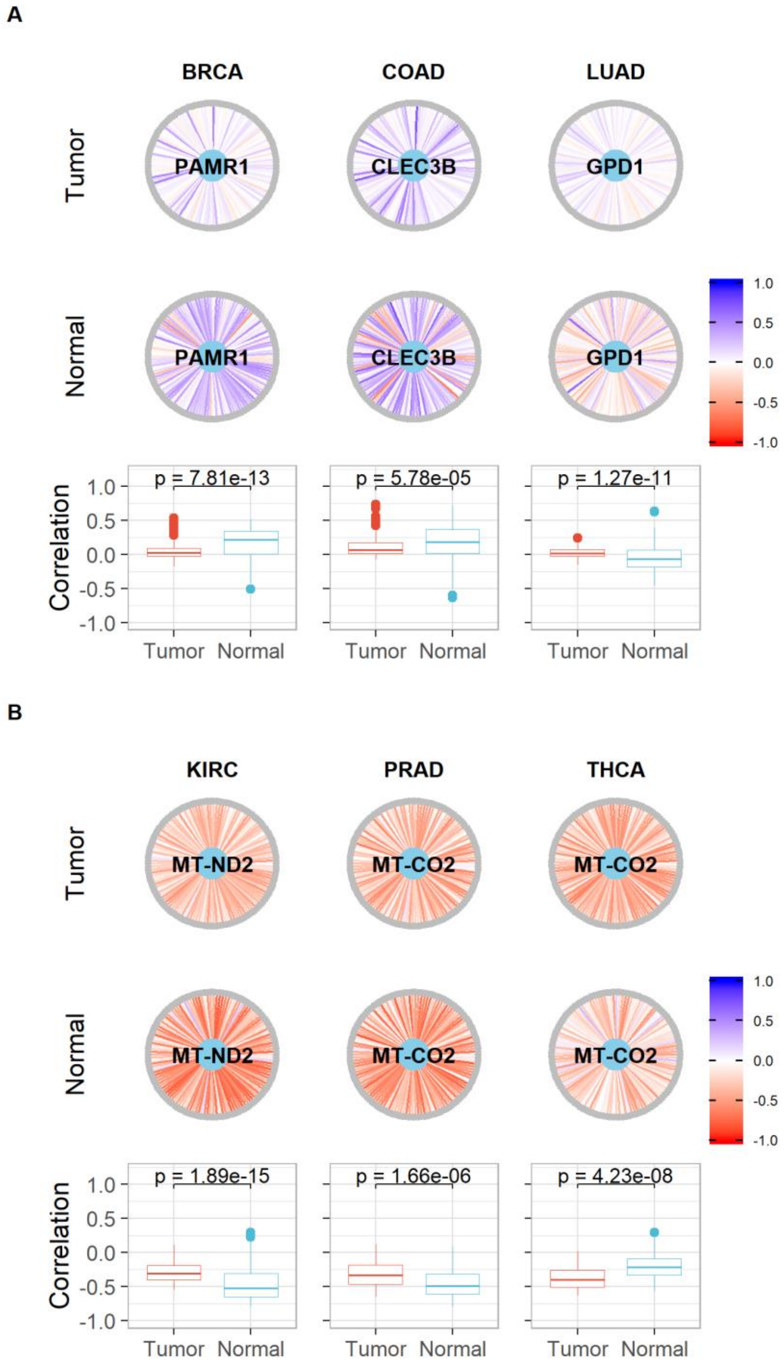

**Supplementary Figure S22 - Staphylococcus aureus infection (hsa05150) and some core gene interactions highlighted by SCOPE**. **A)** Shows differentially expressed core genes (light blue) and their interactions with other genes (grey) in the pathway while **B)** shows similar interactions with non-differentially expressed core genes. Correlations are indicated as edges ranging from red (-1) to blue (+1). Boxplots show the correlations indicated in the same pathways to highlight differences in distributions along with the p-value for the Kolmogorov-Smirnov test with the null hypothesis being that the two samples (correlations of Tumour and Normal tissues) were drawn from the same distribution.

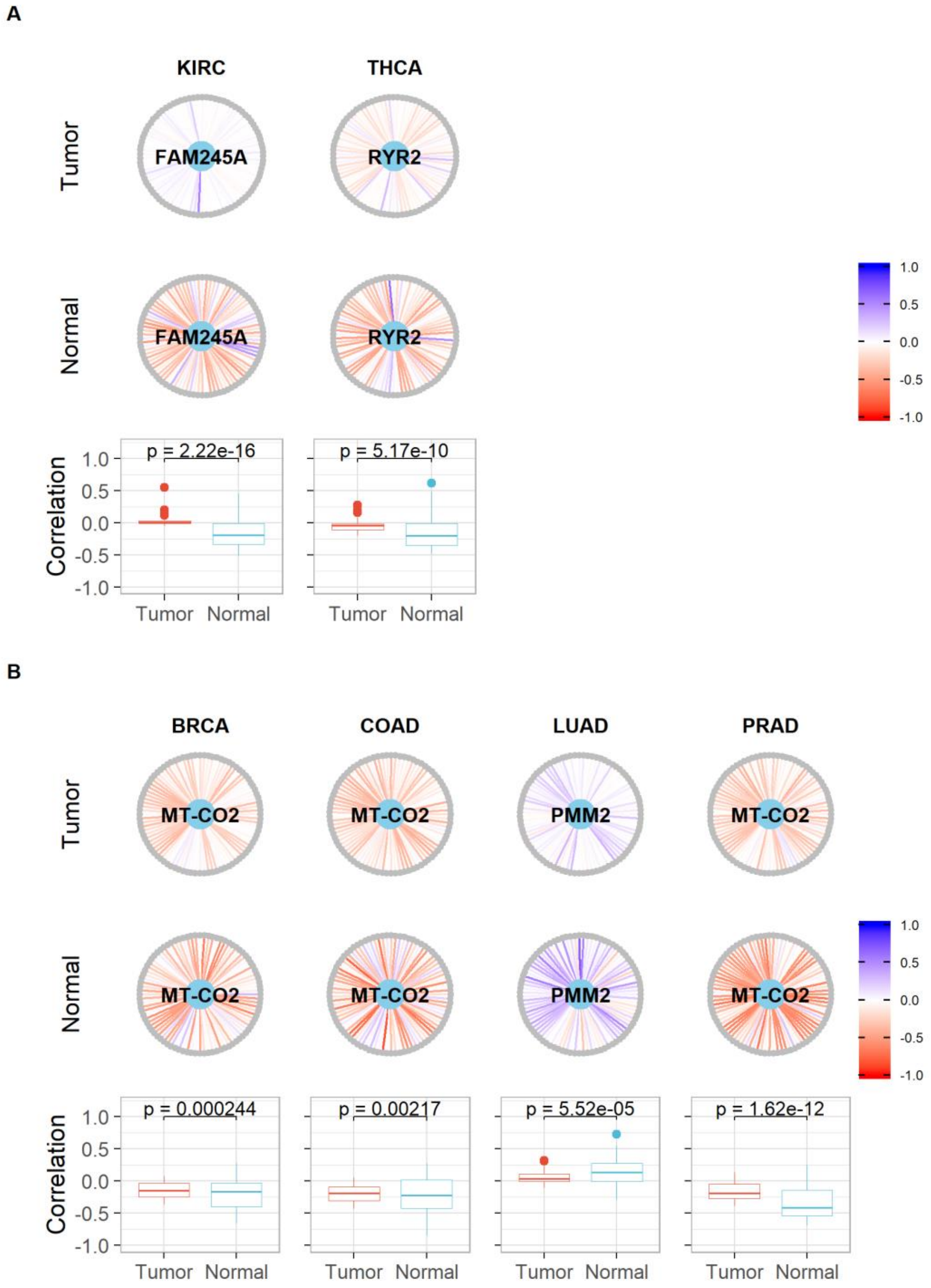

**Supplementary Figure S23 – Network plots of cancer specific pathways.** Core genes are indicated in light blue and other genes in the pathway, in grey. Correlations are indicated as edges ranging from red (-1) to blue (+1).

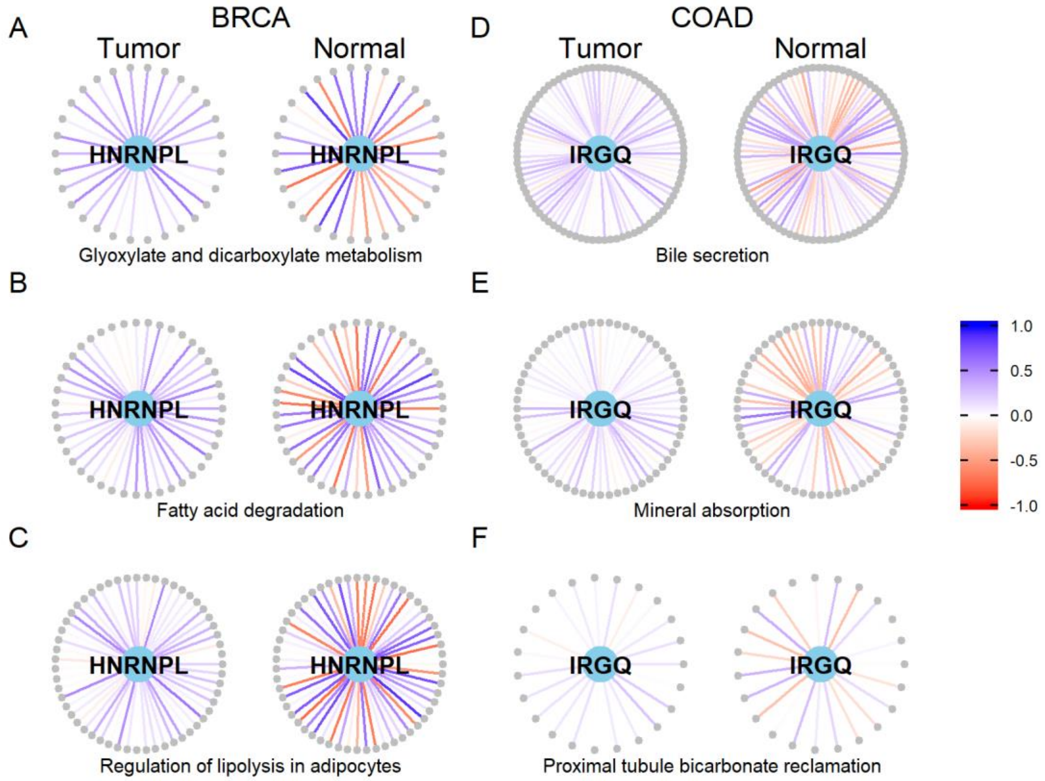

**Supplementary Figure S21 – Survival curves of CD63 over-expressed and under-expressed patients in all cancers.** Log-rank tests indicate a significant difference in survival probabilities only for KIRC with respect to expression levels of CD63. (EXP < 0 indicates samples in which the expression level of the gene (*CD63*) is lower than the arithmetic mean of the expression levels of the gene across all samples; while EXP > 0 indicates higher than mean expression levels.

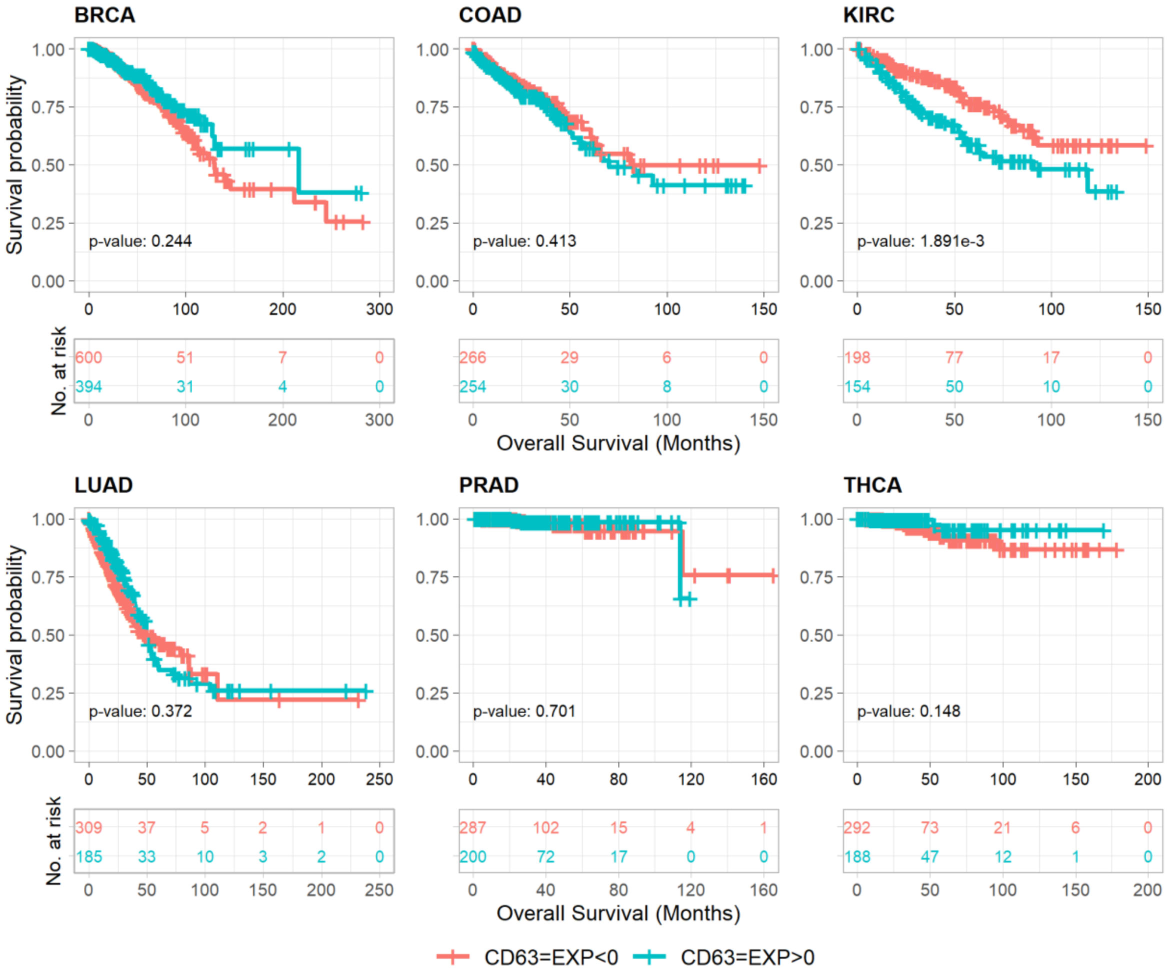

**Supplementary Figure S25 - PI3K-AKT-mTOR signaling pathway** is highly mutated in BRCA. Pathway diagram obtained from cBioPortal.

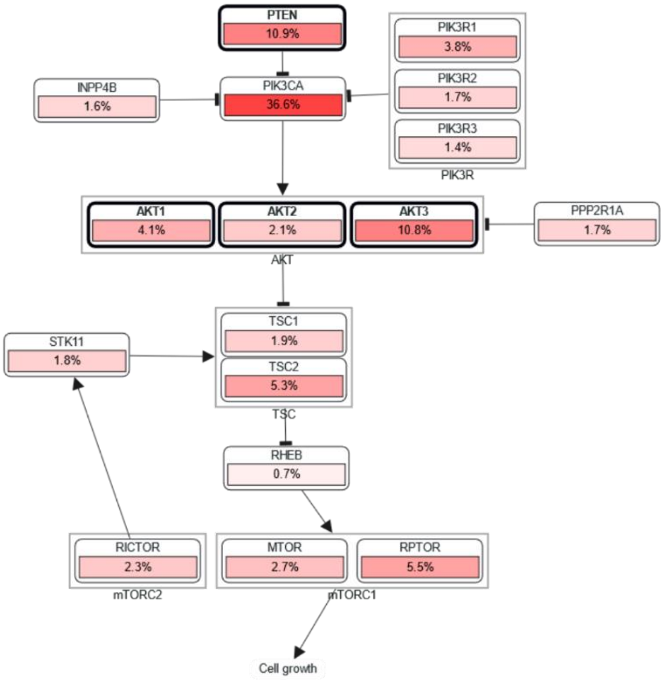
